# Profound structural conservation of chemically cross-linked HIV-1 envelope glycoprotein experimental vaccine antigens

**DOI:** 10.1101/2021.07.26.453798

**Authors:** Gregory Martin, Rebecca A Russell, Philip Mundsperger, Scarlett Harris, Lu Jovanoska, Luiza Farache Trajano, Torben Schiffner, Katalin Fabian, Monica Tolazzi, Gabriella Scarlatti, Leon McFarlane, Hannah Cheeseman, Yoann Aldon, Marielle Breemen, Kwinten Sliepen, Dietmar Katinger, Renate Kunert, Rogier W Sanders, Robin Shattock, Andrew B Ward, Quentin J Sattentau

**Affiliations:** Department of Integrative Structural and Computational Biology, IAVI Neutralizing Antibody Center, Collaboration for AIDS Vaccine Discovery and Center for HIV/AIDS Vaccine Immunology and Immunogen Discovery, The Scripps Research Institute, La Jolla, California, USA; The Sir William Dunn School of Pathology, The University of Oxford, Oxford, UK; Polymun Scientific Immunbiologische Forschung GmbH, Klosterneuburg, Austria; Department of Biotechnology, University of Natural Resources and Life Sciences, Vienna, Austria; Current address: Department of Immunology and Microbial Science, The Scripps Research Institute, La Jolla, California, USA; Department of Immunology, National Food Chain Safety Office, Directorate of Veterinary Medicinal Products, Budapest, Hungary; Viral Evolution and Transmission Unit, Division of Immunology, Transplantation, and Infectious Diseases, IRCCS Ospedale San Raffaele, Milan, Italy; Imperial College London, Department of Medicine, Division of Infectious Diseases, Section of Virology, Norfolk Place, London W2 1PG, UK; Department of Medical Microbiology, Academic Medical Centre University of Amsterdam, Amsterdam, The Netherlands

**Keywords:** HIV-1, envelope glycoprotein, vaccine, chemical cross-linking, structural biology, good manufacturing practice

## Abstract

Chemical cross-linking is used to stabilise protein structure with additional benefits of pathogen and toxin inactivation for vaccine use, but its use is restricted by potential induction of local or global structural distortion. This is of particular importance when the protein in question requires a high degree of structural conservation for the purposes of understanding function, or for inducing a biological outcome such as elicitation of antibodies to conformationally-sensitive epitopes. The HIV-1 envelope glycoprotein (Env) trimer is metastable and shifts between different conformational states, complicating its functional analysis and use as a vaccine antigen. Here we have used the hetero-bifunctional zero-length reagent EDC to cross-link two soluble Env trimers, selected well-folded trimers using an antibody affinity column, and transferred this process to good manufacturing practice (GMP) for clinical trial use. Cross-linking enhanced GMP trimer stability to biophysical and enzyme attack, and had broadly beneficial effects on morphology, antigenicity and immunogenicity. Cryo-EM analysis revealed that cross-linking essentially completely retained overall structure with RMSDs between unmodified and cross-linked Env trimers of 0.4-0.5 Å. Despite this negligible distortion of global trimer structure we identified individual inter-subunit, intra-subunit and intra-protomer cross-links. Thus, EDC cross-linking maintains protein folding, improves stability, and is readily transferred to GMP, consistent with use of this approach in probing protein structure/function relationships and in the design of vaccines.

## Introduction

HIV-1 vaccine design is primarily focused on eliciting neutralizing antibodies (nAb) by targeting the viral envelope glycoproteins (Env), the only target of nAb [1]. Over the past decade a large number of broadly neutralizing antibodies (bnAbs) that target highly conserved surfaces on HIV-1 Env have been isolated from HIV-1-infected individuals, and bnAb infusion into non-human primates and human immune system mice can provide strerilizing immunity [1]. This provides proof of concept that if bnAbs could be elicited by active vaccination, they would be protective. However, Env is metastable and can adopt different conformational states with implications for bnAb binding and elicitation [2, 3]. This is particularly true for soluble forms of Env, which require specific stabilizing mutations to remain in trimeric form. Currently, the two leading approaches to preparing soluble, near-natively folded HIV-1 trimers for vaccine use are either cleaved between gp120 and gp41 with stability maintained by engineered disulfide bonds and other mutations (termed ‘SOSIP’), or cleavage replaced or supplemented by the use of a flexible linker (here termed Uncleaved pre-Fusion Optimized, UFO). Recently a novel SOSIP trimer – ConM - based upon the sequence of all group M isolates, showed native-like morphology by electron microscopy (EM), bound most bnAbs but not non-neutralizing Abs (non-nAbs), and elicited antibodies in rabbits that neutralized pseudoviruses carrying the autologous Env [4]. Similarly, a combined SOSIP - UFO-type design of a group M consensus soluble trimer (ConSOSL.UFO, here termed ConS) was shown by EM to be well-folded in the ‘closed’ conformation and displayed bnAb and non-nAb binding similar to its membrane anchored counterpart [5]. However, despite their relative stability in vitro, these trimers may still sample different conformational states, and their structural and antigenic stability in vivo is unknown, but likely to be detrimentally affected by enzymatic and non-enzymatic modification.

Chemical cross-linking remains widely used for pathogen and toxin inactivation, and for stabilizing proteins for vaccine use and structural analysis. However, how this process affects protein structure at the molecular and atomic level, and how this impacts vaccine antigenicity and immunogenicity is poorly understood and remains empirical for vaccine use. Cross-linking endpoints for vaccine use are generally defined as infectivity reduction for inactivated pathogens [6], or depletion of enzyme activity for toxins [7], with mostly unknown effects on immunogenicity by comparison with the unmodified material. Moreover, we have only a crude understanding of how cross-linking influences vaccine antigenicity. Monoclonal antibody (mAb) binding to hemagglutinin was reduced by formaldehyde (FA)-treatment for inactivated influenza vaccines [8, 9] and to tetanus toxin (TT) to produce the toxoid [10]. Similarly, FA or glutaraldehyde (GLA) treatment reduced pertussis toxin enzymatic and carbohydrate binding activities [11, 12], and FA inactivation of poliovirus inhibited receptor binding [13]. Proteomic and biophysical analyses demonstrated that FA treatment of TT introduced relatively subtle molecular modifications without evidence for major protein conformational changes [14, 15]. However, aldehyde inactivation of vaccines introduces additional atoms from the cross-linking agent, can reduce adaptive immune responses [16], and in some cases such as the prototypic respiratory syncytial virus vaccine, can adversely bias adaptive immune responses driving enhanced disease upon infection [17, 18]. Thus, the use of alternative, more modern cross-linking reagents without these potential adverse effects requires exploration, with the caveat that this process must be Good Manufacturing Practice (GMP) compliant for human use. Moreover, understanding of the structural outcomes of chemically cross-linked vaccine antigens using high-resolution structural analysis is almost completely lacking in the literature, as is the effect of these modifications on the magnitude and specificity of ensuing B and T cell responses. Acquisition of this knowledge is critical, as it would allow prediction of the effects of cross-linking on protein structure, and more rational translation of chemical cross-linking approaches to the design and development of vaccine antigens.

To further stabilize ConM and ConS Env trimers whilst avoiding the known pitfalls of aldehyde cross-linking, we turned our attention to the zero-length heterobifunctional carbodiamide cross-linker 1-Ethyl-3-[3-dimethylaminopropyl]carbodiimide hydrochloride (EDC). This cross-linker activates carboxyl groups on acidic amino acid (aspartic acid, glutamic acid) side chains that then react to form a covalent amide bond with proximal amines on basic amino acid (lysine, arginine) side chains, without addition of any additional atoms to the molecule. To our knowledge this cross-linking reagent has not previously been used to stabilize vaccine antigens. Here we have cross-linked ConM and ConS trimers using EDC, and produced them to GMP for experimental medicine trials. We show that EDC cross-linking enhances trimer stability against denaturation and protease attack, whilst preserving structural integrity to a striking extent. Consistent with the preservation of structure, cross-linking had largely beneficial effects on trimer antigenicity, and immunogenicity in mice and rabbits. Thus, antigen stabilization by EDC cross-linking results in minimum structural and functional perturbation, can be translated into HIV-1 vaccine immunogens for human use, and is an approach generally applicable to other vaccines.

## Results

### Biophysical and antigenic characterization of trimers

ConM and ConS trimers developed and characterized as previously described [4, 5], were EDC (**Fig. 1A**) cross-linked using optimized conditions, which were translated to a GMP workflow as outlined in **Fig. 1B**. In brief, trimers were cross-linked with EDC in a process modified from that previously described for the BG505 trimer [19], and purified using an affinity column made with the PGT145 bnAb manufactured to GMP, to select for well-folded forms. The PGT145 apex-specific bnAb was chosen since it is a stringent probe of trimer conformational integrity [20], and binds very similarly to unmodified and EDC cross-linked ConM and ConS trimers (**Supplementary Fig. 1**). Analysis of unmodified primary amines (**Fig. 1C**) revealed no difference between unmodified and cross-linked ConM, whereas EDC modification of ConS reduced amine content by ∼15%, suggesting more extensive but still relatively modest modification. Cross-linked trimers were resistant to reducing-denaturing SDS-PAGE compared to their unmodified counterparts, which dissociated into their respective monomeric species (**Fig. 1D**). Because the gp120-gp41 subunits are peptide-linked in ConS, they have a higher molecular weight than ConM which dissociates into gp120 and truncated gp41 subunits. As anticipated from the SDS-PAGE outcome, differential scanning caolrimetry (DSC) analysis revealed that cross-linking enhanced thermal stability by 18.7^0^C and 17.6^0^C for ConS and ConM respectively (**Fig. 1E**). Cross-linked ConM yielded an overlapping series of peaks suggesting a heterogeneous population of stabilized trimer forms. Trimer antigenicity was probed by ELISA analysis with panels of bnAbs and non-neutralizing antibodies (non-nAbs - **Fig. 1F and Supplementary Fig. 1**). **Fig. 1F** summarizes the ratio of modified : unmodified trimer binding, such that 1 represents no change, <1 represents reduced binding to modified trimer and >1 represents increased binding to modified trimer. On average, cross-linking reduced bnAb binding by <2-fold and non-nAb binding (with the exception of 19b, excluded from the averaging) almost to background on both trimers. Selective conservation of bnAb binding after cross-linking was observed. Binding to the V3 glycan epitope cluster represented by 2G12 [21] and PGT121 [22] showed no change in binding, whereas to trimer apex (represented by PGT145 [22] and PG16 [23]), CD4bs (VRC01 [24] and 3BNC117 [25]) and interface (35022 [26]) epitope clusters, there was no change or a modest reduction of <2-fold. By contrast a striking reduction in binding was observed upon cross-linking of ConM with the fusion peptide-specific bNAbs ACS202 [27] and VRC34 [28]. ConS does not bind these bnAbs detectably since they preferentially engage the fusion peptide in cleaved trimers [28, 29]. This pattern of bnAb binding to modified and unmodified trimers was confirmed by bilayer interferometry (Octet) analysis (**Supplementary Fig. 2** and **Table 1**). Interestingly, the dramatic loss of VRC34 binding upon EDC modification of ConM was reflected primarily in a highly reduced on-rate (**Supplementary Fig. 2** and **Table 1**), suggesting that bNAb access to its epitope was limited by trimer cross-linking rather than direct EDC alteration of the epitope. This is to be expected since the non-polar fusion peptide epitope for VRC34 does not contain any carboxyl or free amine groups. Non-nAb binding to the cross-linked trimers was essentially absent with the exception of the V3-specific 19b [30] which showed no change after cross-linking (**Fig. 1F**), implying that this epitope is constitutively expressed on all trimer forms regardless of stability. sCD4-IgG2 binding to unmodified trimers was weak, and was abolished by cross-linking, implying that EDC cross-linked trimers will not engage CD4 on T cells, a probable advantage for immunogenicity since CD4 T cell binding could sequester trimers, and particularly the CD4 binding site, from B cell recognition. Similarly, weak binding of the CD4-induced epitope-binding antibody 17b [31] to ConM (17b did not bind ConS) in the presence of sCD4 was eliminated after cross-linking. Two-dimensional classification of trimer morphology by negative stain electron microscopy (EM) revealed that unmodified ConM and ConS trimers were 99% and 78% well-folded respectively prior to modification, and 94% and 99% well-folded subsequent to cross-linking and antibody selection (**Fig. 1G**). Thus, EDC cross-linking and PGT145 affinity column processing may have subtly reduced the proportion of well-folded ConM trimers, whereas the same procedure selected well-folded ConS trimers from amongst a more heterogeneous unmodified population.

**Figure 1.**
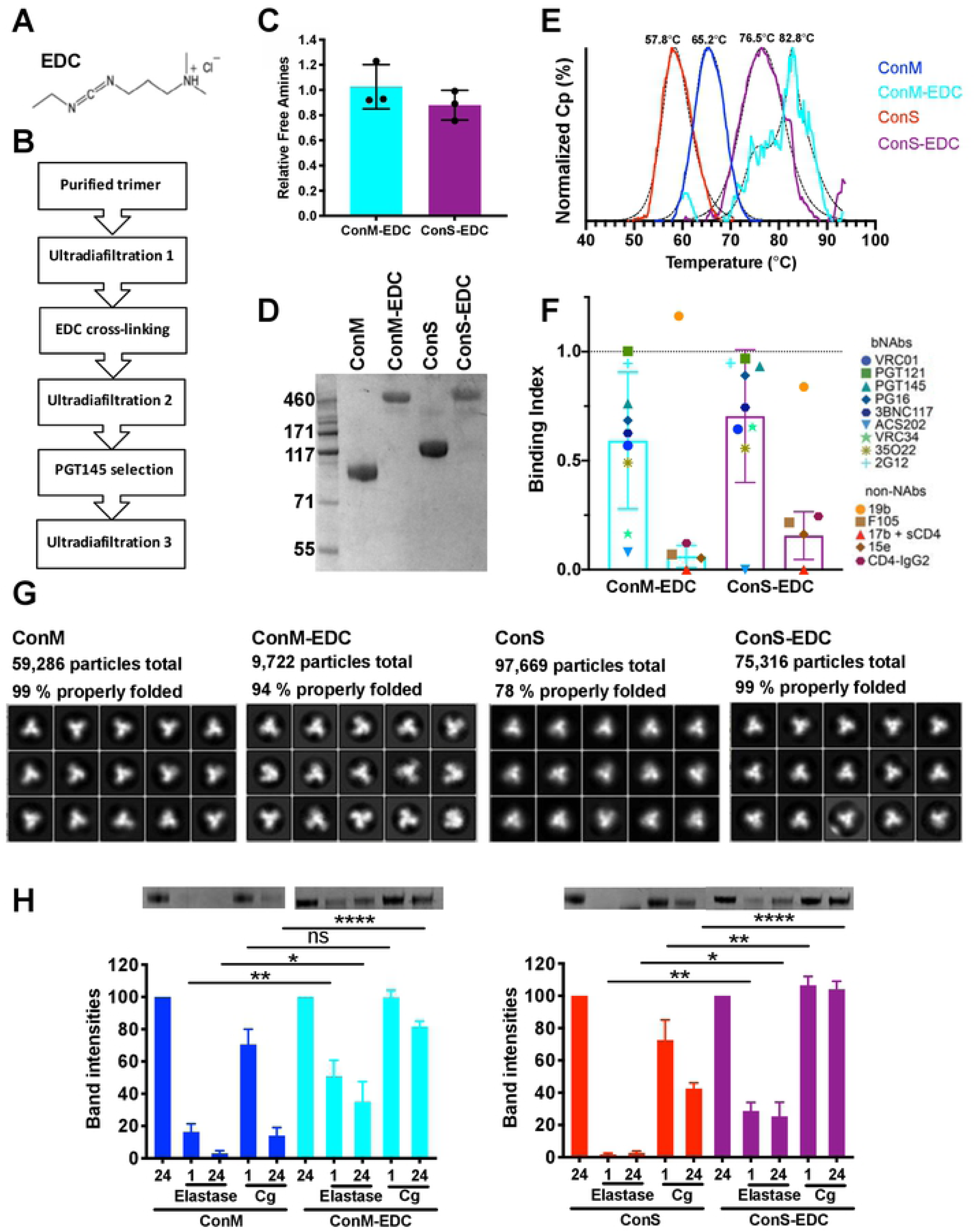
Biophysical and antigenic characterization of GMP EDC cross-linked ConM and ConS. **A**) EDC cross-linker. **B**) Work-flow to GMP product. **C**) Free amine assay representing the ratio of free amines in EDC-modified : unmodified trimer. **D**) SDS-PAGE reducing gel. **E**) DSC analysis of trimer thermal stability. **F**) Antigenicity of trimers using panels of bnAbs and non-nAbs, where results represent ratio of binding modified : unmodified trimer and bars represent ± 1 SD (note 19b excluded from mean). **G**) Negative stain EM 2D classification analyses. **H**) Sensitivity of unmodified and EDC cross-linked trimers to neutrophil elastase and Cathepsin G (Cg) attack, n = 3. ns = not significant; * p<0.05; ** p<0.01; **** p<0.0001, Mann Whitney U.

**Table 1.** Octet analysis of bnAb affinity for unmodified and EDC cross-linked trimers.

| bnAb | trimer | KD (M) | Kon (1/ms) | Koff (1/s) |
| --- | --- | --- | --- | --- |
| 35022 | ConM | $3.9 \times 10^{-7}$ | $4.3 \times 10^3$ | $8.6 \times 10^{-5}$ |
| | ConM-EDC | $2.0 \times 10^{-6}$ | $7.4 \times 10^1$ | $8.3 \times 10^{-5}$ |
| PGT122 | ConM | $<1 \times 10^{-12}$ | $7.3 \times 10^4$ | $<1 \times 10^{-7}$ |
| | ConM-EDC | $<1 \times 10^{-12}$ | $7.8 \times 10^4$ | $<1 \times 10^{-7}$ |
| PGT145 | ConM | $3.0 \times 10^{-8}$ | $1.8 \times 10^4$ | $3.1 \times 10^{-4}$ |
| | ConM-EDC | $1.8 \times 10^{-7}$ | $3.7 \times 10^3$ | $5.1 \times 10^{-4}$ |
| VRC01 | ConM | $1.5 \times 10^{-7}$ | $3.3 \times 10^3$ | $4.2 \times 10^{-4}$ |
|  | ConM-EDC | ND | ND | ND |
| VRC34 | ConM | $1.2 \times 10^{-8}$ | $1.9 \times 10^5$ | $2.2 \times 10^{-3}$ |
| | ConM-EDC | $3.2 \times 10^{-6}$ | $3.1 \times 10^2$ | $2.9 \times 10^{-4}$ |

In animal models, intramuscular administration of adjuvanted vaccine formulations rapidly recruits neutrophils (detectable within 1 h) [32], and subsequently monocytes and macrophages [32–34], to the injection site where they will release bioactive mediators including the major leukocyte proteolytic enzymes elastase and cathepsin G [35]. We therefore hypothesized that adjuvanted protein vaccine immunogens will be subject to localized enzyme attack in vivo. Given the importance of correct folding of Env trimers for promoting the induction of nAb as opposed to non-nAb [36–38], proteolytic cleavage might compromise trimer integrity and thereby limit induction of bnAbs. To test this hypothesis, we exposed trimers to concentrations of elastase (200ng) or cathepsin G (168 ng) equivalent to that estimated to be released by ∼10^5^ and 5×10^6^ neutrophils respectively (data not shown) for 1 or 24 h, and analyzed their integrity by reducing SDS-PAGE. For unmodified ConM trimers this resulted in individual monomer gp120 and truncated gp41 bands, of which the gp120 band was quantified by densitometry, whereas cross-linked trimers ran intact, and these bands were quantified by densitometry. **Fig. 1H** reveals that unmodified ConM trimer was highly susceptible to elastase cleavage, losing >80% integrity within 1 h, and was almost completely hydrolyzed by 24 h, whereas cathepsin G had only modest effects at 1 h, but band density was reduced by ∼85% at 24 h. By contrast, cross-linked ConM showed only ∼50% elastase cleavage at 1 h, significantly less than its unmodified counterpart (p<0.01), reducing to 65% at 24 h, again significantly less than the unmodified trimer (p<0.05). ConS ran as a truncated gp140 band due to the linker peptide between the gp120 and gp41 subunits, and showed high sensitivity to elastase cleavage, with complete band degradation by 1 h, and partial sensitivity to cathepsin G, with ∼30% and ∼60% reduction at 1 and 24 h respectively. As with ConM, cross-linking significantly protected the trimers from degradation under every condition tested (1 h, p<0.001; 24 h, p<0.05), and completely prevented cathepsin attack (1 h p<0.01, 24 h p< 0.0001).

In summary, EDC cross-linking and PGT145 selection of both ConM and ConS trimers enhanced thermal stability and resistance to biophysical and enzymatic denaturation, whilst retaining good trimer morphology and selectively modifying antigenicity in a largely beneficial manner.

### Structural analysis of trimers

Since preservation of HIV-1 Env trimer folding is important for maintaining conformational bnAb epitopes, in particular those dependent on quaternary conformation, it is critical that cross-linking does not adversely impact trimer structure. Moreover, high-resolution structural information relating to cross-linking of highly conformation-dependent proteins would be helpful additional knowledge relating to the use of such stabilizing agents for understanding general principles of protein structure/function relationships. To interrogate the effect of EDC cross-linking we carried out cryo-EM analysis of trimers bound to PGT122 [22] Fab fragments, and solved the structures of unmodified and cross-linked GMP versions of ConS at 3.1 and 3.45Å resolution respectively, and unmodified and cross-linked GMP ConM at 3.4 and 3.85Å respectively (**Fig. 2**, **Supplementary Table 1** and Supplementary **Figs. 3-6**). Each Env is found in the stable closed, prefusion state, and all are highly conserved in overall structure compared to other SOSIP trimers, with an overall Cα Root Mean Square Deviation (RMSD) of 1.25Å (ConS) and 1.2Å (ConM) compared to BG505 (PDB 4TVP; **Supplementary Fig. 7**). The ConS structure is novel, and the ConM cryo-EM structure is similar to the previously published [4] X-ray structure of ConM in complex with PGT124 and 35022 (1.0Å Cα RMSD; PDB 6IEQ), albeit at higher resolution (3.4Å cryo-EM; 3.9Å X-ray). Interaction of PGT122 with ConM and ConS is very similar to other SOSIP trimers, and is driven by contacts with glycans at N332 and N138 (N137 in BG505) and residues at the base of the V3 loop. In agreement with the antigenicity data by ELISA of the related PGT121 bnAb (**Fig. 1F** and **Supplementary Fig. 1**), and by Octet for PGT122 (**Supplementary Fig. 2**), EDC cross-linking has no obvious effect on recognition of these epitopes (**Supplementary Fig. 8**).

**Figure 2.**
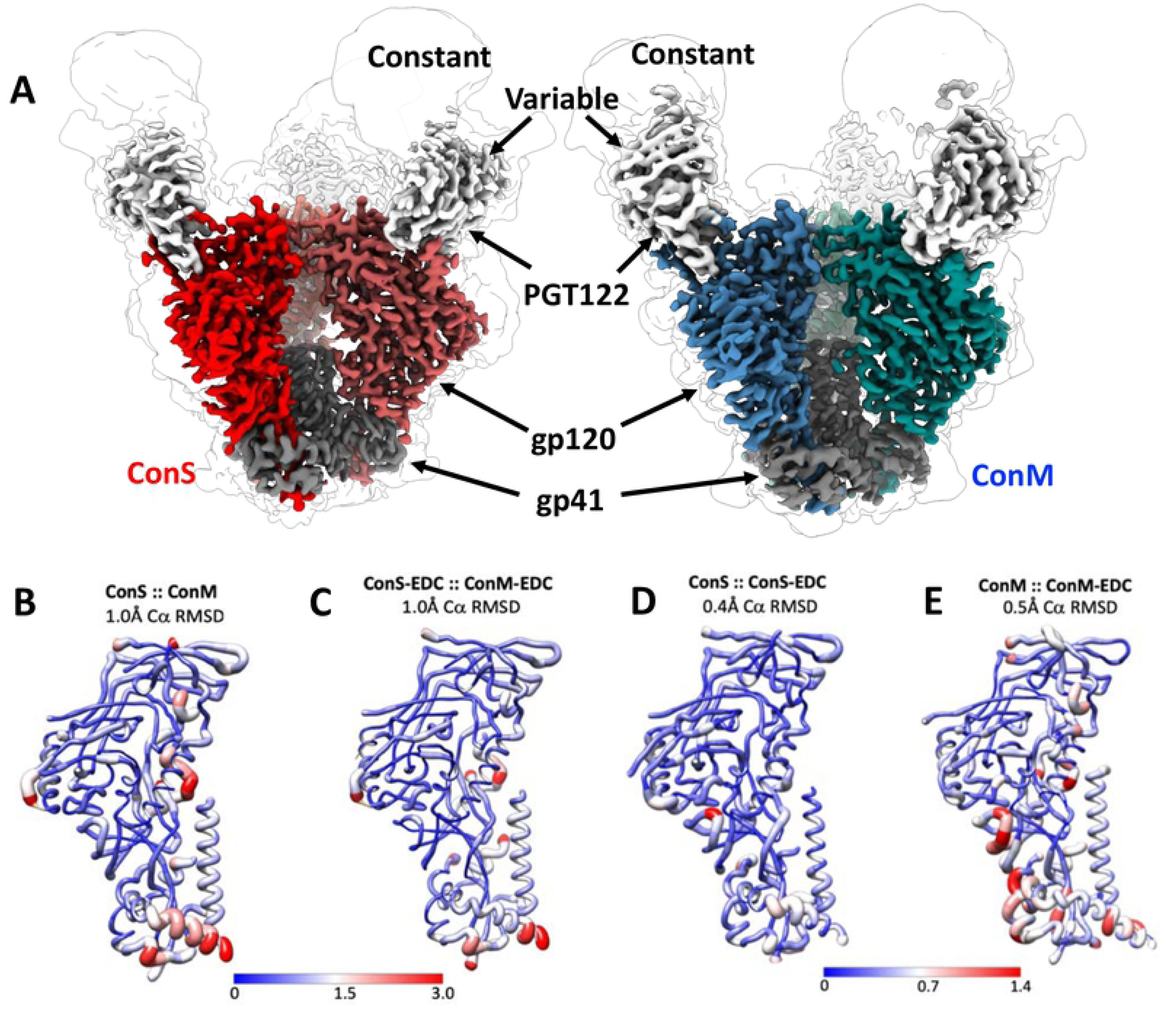
Cryo-EM analysis of ConM, ConM-EDC, ConS and ConS-EDC. **A**) Side views of unmodified ConS (red) and ConM (blue) in complex with PGT122 Fab (grey) at 3.1 Å and 3.4 Å respectively. Each cryo-EM map is shown at high (colored map) and low (light outline) threshold to highlight the PGT122 constant domain and N-linked glycans. (**B)** ConM and ConS structures were superimposed, and local Cα RMSD was rendered onto the ConM structure with Chimera, according to color and to the thickness of the main chain cartoon representation. (**C**) Same as in (**B**) but comparing ConS-EDC and ConM-EDC structures rendered onto the ConS-EDC structure. **(D)** Same as in (**B**) but comparing ConS and ConS-EDC structures, rendered onto the ConS structure. (**E**) Same as in (**B**) but comparing ConM and ConM-EDC rendered onto the ConM structure.

**Figure 3.**
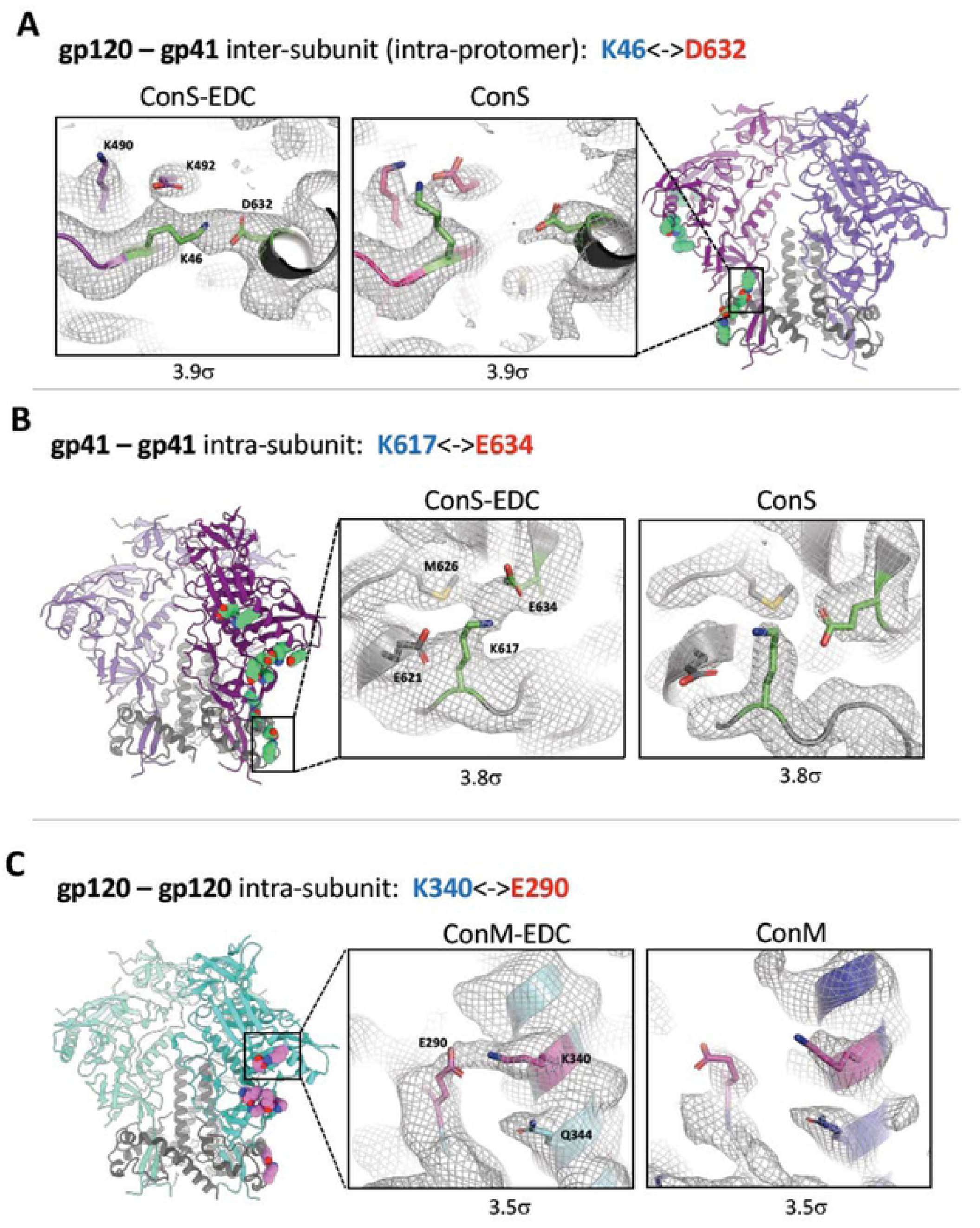
High resolution analysis of EDC-introduced cross-links. **A – C**, selected EDC crosslinks observed through comparison of modified and unmodified cryo-EM reconstructions. Each map is filtered to the same resolution (lower of the two maps) and the same B-factor is applied, and then visualized at the same contour level (σ; see methods). The overall structure is shown in ribbon representation, and the side chains of every crosslink identified are shown as green spheres. **(A)** K46-D632 crosslink in ConS-EDC, which is a gp120-gp41 inter-subunit crosslink within the same protomer. The insets show a zoomed-in view of the K46-D632 crosslink, with cryo-EM density displayed as a mesh. **(B)** Same as in (**A**), for the K617-E634 intra-subunit crosslink in ConS-EDC gp41. Note that the ribbon structure at left is rotated ∼90° counterclockwise (as viewed from the top) from (**A**). **(C)** K340-E290 intra-subunit crosslink in ConM-EDC gp120.

ConM and ConS trimers are both based on consensus sequences of group M isolates, and have ∼91% sequence identity (**Supplementary Fig. 9**) and a Cα RMSD between protomers of 1Å (**Fig. 2B** and **Supplementary Fig. 7**). Inspection of the unmodified ConM and ConS cryo-EM structures identified a number of notable features shared by these two Env designs; among these are the trimer stabilizing mutations A316W and H66R (**Supplementary Fig. S10**). W316 is found at the trimer apex within the V3 loop. In both structures the W316 side chain is highly ordered and mediates a cation-pi interaction with R308, and a hydrophobic stacking interaction with Y318, together stabilizing the β-hairpin structure of the V3 loop and probably impeding V3 remodelling necessary for induction of the CD4 bound state[39, 40]. R66 is found within a loop in the gp120 inner domain, just beneath the apex. Interestingly, in both ConM and ConS maps, we observe density for two rotamers of the R66 side chain, with one conformation exposed to solvent and the other forming electrostatic interactions with T71 and S115. The equilibrium between these two states may underly the mechanism by which H66R inhibits CD4-induced structural changes.

### Structural Impacts of EDC Crosslinking

Since EDC is a ‘zero-length’ cross-linker, we did not anticipate the presence of additional atoms in the cross-linked structures, but the formation of individual amide bonds between proximal carboxyl and amine groups could distort local structure, and multiple cross-links might deform overall protein folding. However, comparison of the unmodified and cross-linked structures revealed a striking near identity, with a Cα RMSD of only 0.45Å between ConS and ConS-EDC, and 0.5Å between ConM and ConM-EDC (**Fig. 2D, E**). Similarly, the RMSD between cross-linked ConM and ConS trimers was also only 1Å (**Fig. 2C**). Thus, within the limits of resolution, EDC cross-linking had a negligible impact on trimer structure. Thus, overall, these data suggest that any impacts to antigenicity and bnAb binding are not a direct result of global structural or conformational changes imparted by EDC crosslinking. Moreover, this highlights a more general point that EDC cross-linking may be used to stabilize conformationally sensitive protein structure without substantial local or global distortion.

Throughout the ConS and ConM structures there are numerous potential EDC crosslinking sites, in which an aspartate or glutamate side chain is in proximity to, or directly interacting with, a lysine side chain (**Supplementary Table 2**). However, as a cryo-EM reconstruction represents an ensemble of many thousands of particles, and in this case contains C3 symmetry, only high-frequency events will be observed in the final map. Strikingly, close inspection of the ConS-EDC and ConM-EDC density maps, and careful comparison with their unmodified counterparts, revealed a series of high confidence cross-links. Observed cross-links are listed in **Supplementary Table 2**, and examples are summarized visually in **Fig. 3**. Most were intra-subunit crosslinks within gp120, with an inter-subunit gp120-gp41 (intra-protomer, **Fig. 3A**) and an intra-subunit within gp41 (**Fig. 3B**). Inter-subunit (gp120-gp41) crosslinks such as K46-D632 (**Fig. 3A**) will stabilize quaternary structure of the complex consistent with the SDS-PAGE analysis (**Fig. 1D**). This should increase stability and in vivo lifetime of the ‘closed’ state of the trimer, thus maintaining bnAb epitopes but reducing exposure of non-nAb epitopes such as the V1V2 and V3 loops and the CD4i surface [19, 41–43]. Since each crosslink is in essence a “molecular staple”, even intra-subunit cross-links (**Fig. 3B, C**) will act to stabilize secondary and tertiary structure, increasing overall molecular stability and potentially reducing exposure to protease attack.

Structural analysis informs antibody binding, allowing comparison between unmodified and cross-linked antigens. Modification of antigenicity may result from global changes in trimer folding, which we have shown is not the case. Thus, changes will most probably be local, and will result from introduction of cross-links within or proximal to the epitope, and potentially also the change in charge at the site of cross-linking arising from amide bond formation that eliminates the previous negative charge of the reacted carboxylate group [44]. However, inspection of the epitope cluster most heavily antigenically modified in ConM - the fusion peptide - (**Fig. 1F, Supplementary Figs. 1 and 2 and Table 1**), revealed no obvious cross-links. However, E87 in gp120 that is reported to be critical for ACS202 binding [29] may be modified by EDC in ConM, reducing bnAb engagement. However, this would not explain loss of VRC34 binding to the same region of ConM, as this bnAb does not include E87 in its epitope[28]. Therefore, it seems most likely that fusion peptide exposure is prevented by the trimer cross-linking process. By contrast, whilst none of the CD4bs, apex and interface epitopes analyzed contained obvious cross-links, all contained acidic residues that could form heterogenous low-frequency, unobserved crosslinks with proximal lysines (**Supplementary Fig. 11**), potentially resulting in reduced affinity which would be consistent with the modified binding curves (**Supplementary Fig. 2**) and reduced k_D_ values (**Table 1**). An alternative is that the quenching step to eliminate unreacted O-acylisourea intermediates in the absence of proximal primary amines might form an amide bond with quenching agent (glycine), the added glycine residues disrupting antibody epitopes. However, this seems unlikely since no extra density was observed on aspartate or glutamate side chains (data not shown), suggesting that if it does occur, it is likely to be a rare event. Moreover, as mentioned previously, EDC cross-linking primarily influenced bnAb on-rate, strongly implying reduced epitope exposure rather than direct impact on epitope chemistry.

Taken together, these data demonstrate a striking conservation of structure in cross-linked trimers with highly localized and epitope-specific modulation of bNAb binding evidenced by high-resolution structural analysis.

### Adaptive immune responses to trimer immunization

Modification of antigenicity by cross-linking may influence trimer immunogenicity with respect to adaptive immune recognition of B and T cell epitopes. Preclinical analysis is of particular importance for GMP material as it will help to predict immune responses in experimental medicine trials. To probe antigen-specific adaptive immune responses, mice were immunized with 10 µg trimer/dose formulated in 10 µg/dose MPLA adjuvant at the times indicated by the arrows (**Fig. 4A, B**), and antibody kinetics analysed by ELISA. Serum from each group was assayed for antigen-specific IgG against either unmodified or EDC cross-linked homologous trimers at the times shown. Unmodified ConM elicited an IgG response that reacted equivalently with unmodified and cross-linked ConM at week 2, but which then diverged after the prime showing significantly greater reactivity against unmodified compared to cross-linked ConM at 6 weeks (p<0.0001) and 12 weeks (p<0.01), **Figure 4A**. Similar results were obtained for ConS (**Fig. 4B**), with sera from ConS-immunized mice giving significantly higher titres on ConS compared to EDC cross-linked ConS at week 6 (p<0.01). These data suggest that epitopes available on unmodified trimers, potentially as a result of trimer opening and/or dissociation in vivo, are less represented on cross-linked trimers, which are unable to open and dissociate into their components. By contrast, sera from mice immunized with cross-linked ConM and ConS trimers gave very similar titres on both unmodified and cross-linked homologous trimers, consistent with epitopes available on the cross-linked trimers being equivalently presented on the unmodified trimers.

**Figure 4.**
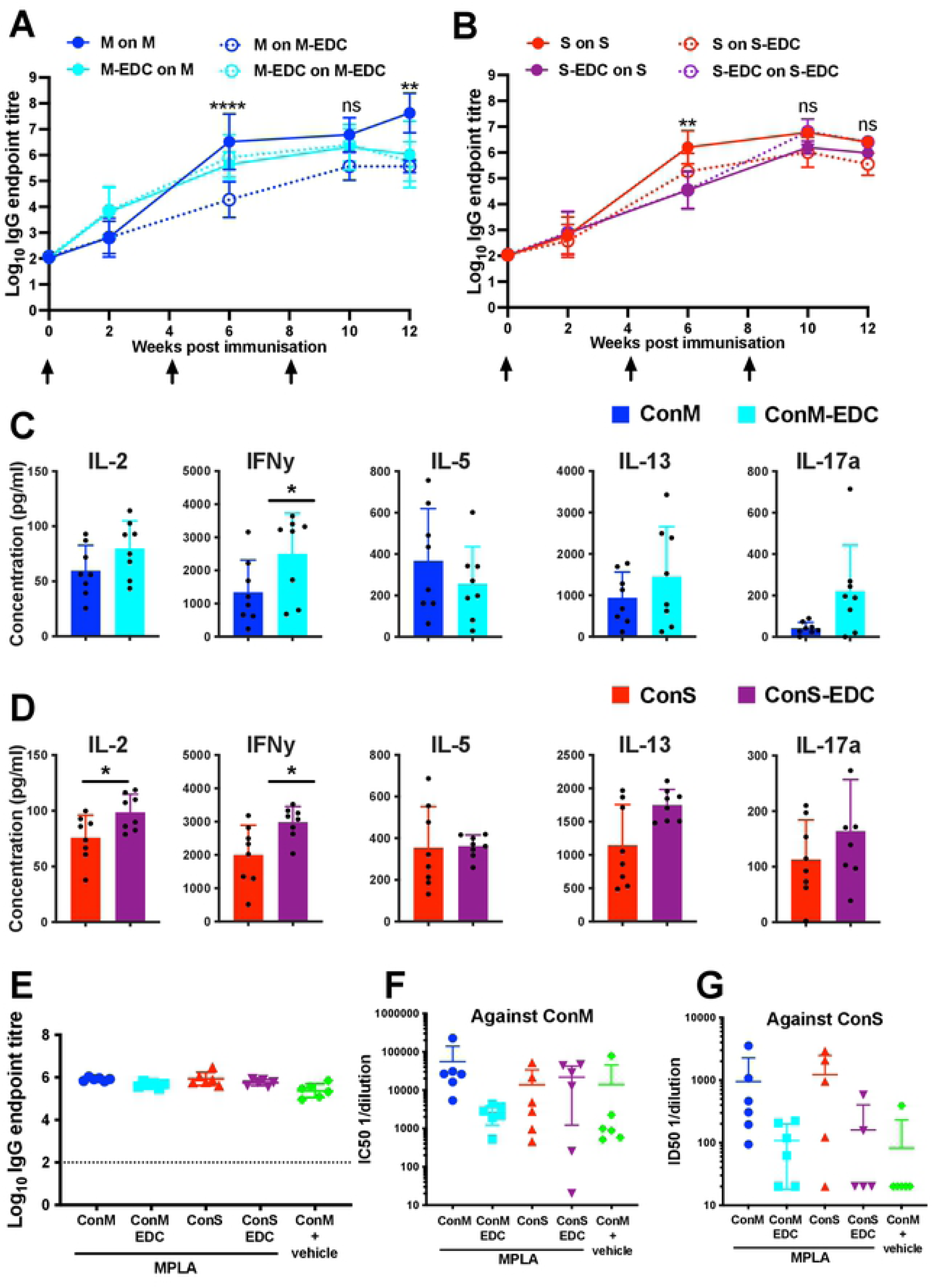
Immunogenicity of unmodified and cross-linked trimers. **A, B**) Mouse serum IgG responses to immunization with (**A**) ConM (M) or ConM-EDC (M-EDC), or (**B**) ConS (S) or ConS-EDC (S-EDC), assayed against unmodified (M or S) or crosslinked (M-EDC or S-EDC) versions of the protein, where n= 8 mice, comprised of 4 mice/group from 2 pooled independent experiments. **p<0.01, ****p<0.001, One-way ANOVA with Sidak’s multiple comparison correction. **C, D**) Mouse splenocyte antigen-specific cytokine release measured by multiplex bead array after unmodified ConM (**C**) or ConS (**D**) in vitro restimulation with homologous unmodified antigen. **E**) Endpoint ELISA titers of rabbit antiserum at week 12. **F, G**) Neutralization activity against homologous and heterologous ConM (**F**) and ConS (**G**) env pseudoviruses represented by the reciprocal serum dilution giving 50% inhibition (IC_50_). *p<0.05, **p<0.01, ****p<0.0001, Mann Whitney U.

Since cross-linking might modify antigen processing and peptide presentation to T helper (Th) cells, we evaluated antigen-specific responses to immunization. Spleens were harvested at week 14 and splenocytes restimulated in vitro with unmodified homologous antigen for 3 days, followed by analysis of cytokine release. In vitro ConM restimulated splenocytes from mice immunized with unmodified or cross-linked ConM trimer showed a non-significant trend towards enhanced IL-2, IL-13 and IL-17a but a significant (p<0.05) increase in IFNγ release in cross-linked compared to unmodified trimer-immunized mice (**Fig. 4C**). Similarly, restimulation of splenocytes from mice immunized with cross-linked ConS trimer resulted in significantly increased IL-2 and IFNγ (p<0.05) and a trend towards increased IL13 and IL17a release. Thus we anticipate that immunization with cross-linked trimer may elicit a modest enhancement of Th cell responses that appear to be balanced between Th1,Th2 and Th17.

Toxicity analysis of the GMP product required immunization of rabbits with trimer (100 µg) co-formulated with MPLA adjuvant (500 µg), according to the regimen in **Supplementary Fig. 12A, B**. IgG purified from rabbit sera were titrated onto ELISA wells coated with the antigens shown, and titration curves generated (**Supplementary Fig. 12C-F**). Binding curves showed no significant differences between groups in kinetics of IgG elicitation, and endpoint titers determined at week 12 were almost identical between all adjuvanted groups (**Fig. 4E**). Thus, differences noted with the mouse immunizations were not recapitulated in the rabbits, potentially because of the much higher (20-fold) dose of antigen used may have masked differences. Week 12 rabbit sera were assayed against infectious molecular clones (IMCs) expressing Envs from ConM and ConS in a TZMbl assay, and results expressed as reciprocal serum dilution yielding 50% inhibition (IC_50_). Sera from rabbits immunized with unmodified ConM neutralized ConM (**Fig. 4F**) and ConS (**Fig. 4G**) IMCs approximately 30-fold and 10-fold more respectively than sera from animals that received cross-linked ConM, although the lack of statistical significance mean that this was only a trend. Similarly, cross-linked ConS trimer showed reduced autologous neutralization on ConS IMC, but again this was not significant. Since both ConM and ConS Envs generate relatively neutralization sensitive Tier-1-type pseudoviruses [4, 5], most of the serum neutralization activity observed is likely to be against the V1V2 loops, as previously described [4]. Therefore, the reduction in serum neutralization probably represents either beneficial masking of these structures by cross-linking V1V2 within the trimer context, or local EDC modification of the specific epitopes recognized by these Tier-1 virus neutralizing antibodies. We suspect that the first hypothesis is the more likely as the animals will mount a polyclonal response to these epitopes, and so epitope-specific modification seems less likely than domain masking. Heterologous Tier-2 virus neutralization was not tested since the short immunization regimen was designed as a toxicity study and was not optimized to elicit nAb, and so we did not anticipate broadly neutralizing responses.

In summary, immunization of mice and rabbits with GMP EDC cross-linked trimers formulated in MPLA adjuvant revealed broadly similar kinetics of antigen-specific IgG induction in mice and rabbits between unmodified and cross-linked trimers, with a modestly enhanced Th cytokine response in mice against the cross-linked trimers. Trimer cross-linking reduced induction of autologous virus neutralization in rabbit sera potentially via masking of V1V2 epitopes.

## Discussion

The structural analysis of complex antigens has been facilitated over the past decade by the use of cryo-EM single particle analysis, allowing for near-atomic resolution. This has led to determination of a variety of structures such as those here, which provide a detailed snapshot of the effects of chemical cross-linking on two GMP experimental vaccine immunogens destined for clinical use. The most striking outcome from the present analysis is the almost complete conservation of trimer structural integrity after robust trimer stabilization by cross-linking with EDC, which was unexpected. Consistent with the structural conservation is the linked broad conservation of antigenicity, with the only major loss of bnAb binding occurring to the fusion peptide in ConM trimers. Whilst ACS202 binding may potentially be compromised by EDC modification of E87, the reason for VRC34 binding is currently unclear as its epitope does not contain charged residues [28]. Aside from the loss of fusion peptide bnAb binding, we anticipate from the antigenic analysis of cross-linked trimers that other major bnAb epitope clusters should be well conserved, and therefore have the potential to trigger bnAb-reactive B cells. To our knowledge only one other study has investigated the structure of a cross-linked vaccine immunogen compared to its un-crosslinked counterpart at high resolution [43]. In this previous study we showed that glutaraldehyde, a 5 carbon homobifunctional amine cross-linker, introduced cross-links into the prototypic SOSIP trimer BG505, which subtly modified global molecular conformation, and eliminated bnAb binding to quaternary epitopes at the trimer apex [43]. However, that study revealed that GLA added extra atoms to the structure with the added risk of creation of immunogenic neoepitopes, and moreover did not analyze GMP produced material and hence is distal from translation to human use.

Our current results reveal that chemical cross-linking of two M group consensus sequence-derived Env trimers substantially enhanced biophysical stability. It is widely accepted that enhancing stability of subunit immunogens, particularly those with metastable characteristics such as HIV-1 Env, is likely to increase in vivo lifetime and reduce exposure of immunodominant non-neutralizing epitopes such as the V1, V2 and V3 loops [39, 40, 45, 46]. In this respect the inter-subunit cross-link K46-D632 may be particularly relevant since as it is likely to substantially restrict molecular movement. However, it is likely that intra-subunit cross-links and modifications will also play a role in restricting molecular movement via secondary and tertiary structure stabilization. Additional benefits of stabilization may also include increased in vivo lifetime, particularly in the presence of antibodies that trigger disassembly of non-cross-linked trimers [47], and resistance to proteases found at sites of inflammation such as at vaccine injection sites. The local in vivo milieu after administration of vaccine formulations has not been well studied, but is likely to be hostile to proteins in terms of proteolytic attack, particularly when the immunogen has stimulated an inflammatory environment containing cells such as neutrophils that release highly active proteases such as elastase [32–34]. Our in vitro models of protease attack revealed that unmodified trimers were sensitive to cleavage by elastase, and to a lesser extent by cathepsin G, but that cross-linking significantly enhanced protease resistance. Measurement of antigen protease sensitivity over 1 and 24 h seems relevant, since a recent study suggests that antigen may persist at the site of intramuscular administration for several days in an adjuvant-dependent manner, and its elimination can substantially reduce ensuing adaptive immune responses [34]. Our demonstration of enhanced trimer resistance to leukocyte elastase and cathepsin G in vitro may have in vivo significance that has not so far been considered. Future studies will be aimed at characterizing the in vivo fate of conformationally-sensitive immunogens such as soluble Env trimers by the use of highly conformation-dependent bNAb probes.

Whilst the minimal modification of trimer structure and antigenicity upon cross-linking suggests that immunogenicity may not be substantially affected, the complexity of the immune system is such that this is impossible to predict. We therefore carried out immunization of mice with the unmodified and cross-linked trimers formulated with the TLR4 agonist adjuvant MPLA. Interestingly, quantitative differences were seen in mouse T cell cytokine responses, with significantly increased titres of IFNγ (ConM-EDC and ConS-EDC) and IL-2 (ConS-EDC), compared to unmodified trimer, suggesting enhanced Th responses. It is unclear why cross-linking might elicit these increases in cytokine response, particularly in light of the minimal structural changes observed, however cross-links and other EDC modifications made to acidic residues may alter antigen processing and presentation to T cells resulting in cytokine modulation. These modulated cytokine responses did not appear to result in any obvious differences in outcome for B cells, given the similar antigen-specific IgG responses. Small differences in antigen-specific IgG responses were observed in mice, but these normalized between the unmodified and cross-linked groups after the first (ConM) and second (ConS) boosts respectively. Rabbit immunizations were designed to test formulation toxicity and so were not optimized for eliciting neutralizing antibody responses. Nevertheless, autologous and heterologous Tier-1a (ConM) and Tier-1b (ConS) pseudovirus neutralizing responses were observed. Interestingly both autologous (in ConM and ConS sera) and heterologous (in ConM sera) responses tended to be reduced by cross-linking, suggesting that one or more Tier-1 type neutralization epitopes was being masked or otherwise compromised. Since one of the goals of immunogen design is to focus immune responses away from regions that may compete for production of bNAbs, such reductions in Tier-1 neutralization may be helpful. Future studies will address the target specificity of these responses and the mechanism of their reduction in cross-linked trimers.

The increasing ease with which complex molecules such as the HIV-1 Env trimer can be structurally analysed by cryo-EM allows the unprecedented high-resolution analysis of post-translational modifications such as chemical cross-linking, which up to now have not been interrogated in vaccine antigens. This information will enable a more rational approach to immunogen stabilization, particularly for molecules that are structurally and/or conformationally metastable. Equally, the proof of concept shown here for the use of EDC in manufacturing a GMP immunogen, currently in experimental medicine trials, paves the way for use of this, and other newer cross-linking reagents, in vaccine manufacture. Indeed, the EDC process added only two additional steps to the GMP process. Since EDC adds no additional atoms to the cross-links, unlike for example glutaraldehyde, there will be minimal risk of generation of neoepitopes with associated concerns of antigenic modification potentially associated with cross-reactivity and autoimmunity.

## Materials and methods

### Antibodies and ligands

Antibodies VRC01 [24], PGT145 [48], 2G12 [49], PGT122 [48], 19b [30], 35022 [26], ACS202 [27] and VRC34 [28] were expressed in freestyle 293F cells under serum-free conditions and purified by protein A chromatography (Thermofisher) following manufacturer’s instructions. Soluble CD4 (sCD4) [50], CD4-IgG2 [51], 15e [30], F105 [30], 17b [31], 19b and PG16 [52] were from the IAVI Neutralizing Antibody Consortium. Fragments antibody-binding (Fabs) VRC01, PGT122, PGT145, and 35022, used in BLI-Octet studies, were expressed in Freestyle 293F cells under serum-free conditions and purified as previously described (Rantalainen et al. 2020. *Cell Reports*). VRC43 Fab was a kind gift from P Kwong and the VRC. Where required antibodies were biotinylated using EZ-link NHS-LC-Biotin according to the manufacturer’s instructions (Thermofisher), or were attached to cyanogen bromide-activated agarose using the manufacturer’s protocol (GE Healthcare).

### Generation of CHO cell lines for stable expression of ConM and ConS Env trimers

Stable Chinese Hamster Ovary (CHO) cell lines expressing ConM and ConS Env trimers were generated by transfection of a parental CHO K1 host cell line using BAC (Bacterial Artificial Chromosome) vector technology [53]. The transfection strategy for ConM (but not ConS) involved co-transfection (ratio 1:1) of a second BAC vector harbouring the gene coding for the human pro-protein convertase (pc) furin for efficient cleavage of the Env precursor in the trans-Golgi network (TGN) [54]. Positive transfected cell pools for both trimers were selected in serum-free medium (CD CHO (Gibco), supplemented with selection medium (SM, 8 mM L-glutamine and phenol red, with 0.5 mg/mL G-418, Sigma). Cell pools were maintained at 37°C, 5% CO_2_ and 80% humidity in a shaking incubator (Kuhner) at 125 rpm, and regularly checked for onset of cell growth until cell viability reached >95 % with steady growth. ConM and ConS expressing cell pools were expanded to 125 mL shake-flasks (Corning) and were split every 3 to 4 days to starting cell concentrations of 2-3 x 10^5^ cells x mL^-1^, in SM). Cell concentrations and cell viabilities were monitored using a Bioprofile CDV cell counter (Nova biomedical, US) and a Multisizer 4e (Beckman Coulter, US) instruments. At each passage supernatant samples were collected for quantification of ConM and ConS Env titres by PGT145 binding ELISA.

#### Isolation of ConM and ConS Env trimer clonal lines

Clonal lines of ConM and ConS transfected cell pools were obtained through limiting-dilution single-cell-cloning. Cell pools were diluted to a limiting cell concentration of 1 cell per 40 µL well aliquot in conditioned single-cell-cloning medium (1:1) mixture of SM, and 0.2 µm filtered CHO medium from a 3-day passaging harvest from cultivation of the parental CHO K1 host cell line, supplemented with rhAlbumin (Sigma Aldrich), rEGF (Repligen) and rTransferrin (Merck). Diluted cell suspensions were plated into 384-well plates (Corning), centrifuged at 200 x g for 5 min, and imaged with a CellMetric (Solentim) imaging device to capture single-cell plating events, and to follow outgrowth of clonal cell populations. Clonal ConM and ConS cell populations were then expanded to 96-well plates (Thermo Scientific) and evaluated for cell growth using the CellMetric imaging system, and expression of ConM and ConS Env trimers was measured by PGT145 ELISA at the end-point of two consecutive passages. Based on expression levels and cell growth 20 ConM and ConS lead clones were selected, expanded into 25 cm² tissue culture flasks (Greiner) and cultivated for three consecutive passages. Sampling at each passage end-point (every 3^rd^ to 4^th^ day) included measurement of cell concentration and viability. Further, cell-free supernatant was collected for analysis of Env trimer expression by PGT145 ELISA. Additionally, cell pellets of the 20 ConM lead clones were analyzed for the co-transfected hFurin construct by standard end-point PCR and hfurin specific primers. Based on cell growth and PGT145 ELISA expression data, ConM and ConS Env cell-specific productivity was calculated on the basis of which lead clone number was further reduced to eight, which were then cryo-preserved and stored in liquid nitrogen.

#### Evaluation of ConM and ConS Env trimer expressing CHO lead clone candidates

In order to identify optimal clones to enter the GMP manufacturing pipeline, small-scale shake-flask fed-batch experiments were performed. Fed-batches were performed in ActiCHO P medium (GE Healthcare) supplemented with 8 mM L-glutamine and phenol red in 125 mL shake flasks (Corning), at a working volume of 45 mL and a seeding cell density of 0.3 × 10^6^ cells × mL^-1^ and a target stop criterion of culture viabilities ≤ 80%. Addition of feed media, ActiCHO Feed A and ActiCHO Feed B (both GE Healthcare) started on day four based on a glucose-controlled feeding regimen. Feed A was added in order to meet a target glucose concentration of 6.5 g × L^-1^. Feed B was added as a constant 0.28% (v/v) feed based on the actual culture volume. Sampling of fed-batch cultures for monitoring of cell concentrations and culture viability, as well as HIV Env trimer concentration and levels of glucose, lactate, ammonium, L-glutamine and L-glutamate were performed on the day of seeding, and continued on day 4 until the end of the process (culture viability ≤ 80%). Further, ConM and ConS lead clones were monitored for their ability to maintain cell specific growth rates [µ] and specific productivities [q_p_] of trimer expression over a period of ≥70 population doublings (PDL) under routine cultivation conditions. Following small-scale fed-batch evaluation, stability monitoring, and selection of the final ConM and ConS lead clones to enter the GMP pipeline, master cell banks (MCBs) were prepared and cryo-preserved in liquid nitrogen under GMP conditions.

#### Scale-up of ConM and ConS expressing CHO lead clones for GMP manufacture

Scale-up and process development for trimer production in fed-batch mode was performed in a ReadyToProcess WAVE 25 bioreactor system using ActiCHO P medium (GE Healthcare) and the respective feed media ActiCHO Feed a and ActiCHO Feed B (both GE Healthcare). Inoculation cultures were obtained by expansion of ConM and ConS lead clones from MCBs in production medium (ActiCHO P medium supplemented with 8 mM L-glutamine) in 1000 mL shake flasks (Corning) in a shaking incubator (Kuhner) (125 rpm, 37 °C, 5% CO2, 80% relative humidity) to support a starting cell density of 0.3 × 10^6^ cells × mL^-1^ in a starting volume of 8 L. WAVE 25 bioreactor runs were operated at 37°C, dissolved oxygen was set to 30%, with no additional pH control by base addition. Sampling of fed-batch cultures was performed in 24-h intervals for monitoring of cell concentration, cell viability, levels of glucose, lactate, ammonium, l-glutamine and l-glutamate. Additionally, cell-free supernatant was collected to measure product concentration by PGT145 ELISA. Feeding of WAVE fed-batch cultures was performed as previously outlined for small-scale fed-batches experiments. At harvest, cells were removed by centrifugation and cell-free supernatants were passed through a positively charged 3M™Zeta Plus™ depth filter (3M) for removal of negatively-charged contaminants. As a final step, depth filtered supernatants were sterile filtered through a Sartopore 2 (Sartorius) filter unit and stored until further processing at 2-8 °C.

#### GMP production of ConM and ConS Env trimers in fed-batch mode

GMP manufacture of ConM and ConS Env trimers was based on previously established master cell banks of ConM and ConS expressing lead clones and cultivation in fed-batch mode using a ReadyToProcess WAVE50 Bioreactor system (GE Healthcare) and purification of well-folded Env trimers from harvested culture supernatant by PGT145 affinity chromatography and subsequent polishing steps. In a nutshell, inoculum cultures were prepared from cryo-preserved master cell banks and cultures were expanded in production medium (ActiCHO P medium, supplemented with 8 mM L-glutamine) to support starting cell densities of ≥ 3 x 10^5^ cells x mL^-1^ at a starting volume of 17 L. WAVE GMP fed-batch production runs were operated at 37 °C, pH was set to 7.0 (controlled via base addition and purging with CO_2_), 20 rpm rocking speed (6° angle) and 30% dissolved oxygen (DO). Daily sampling was performed for analysis of cell concentration, viability, ConM and ConS Env concentration, glucose levels and key metabolites. Addition of nutrient feeds ActiCHO Feed A and ActiCHO Feed B (both GE Healthcare) started when glucose levels dropped below ≤ 4 g x L^-1^ and was continued daily thereafter. ActiCHO Feed A was added to a final glucose concentration of 6.5 g x L^-1^, ActiCHO Feed B was added as a 0.28% (v/v) volume feed based on the working volume in the bioreactor. Once viabilities dropped below 80% (≥ 60%) fed-batch cultures were terminated and culture supernatants were harvested by 2-stage depth filtration using 3M™ ZetaPlus™ filters for cell removal, followed by removal of host cell proteins (HCP) and DNA by anion-exchange (AEX) membrane absorption using 3M™ Emphaze™ Hybrid Purifier filters, followed by a final 0.2 µm sterile filtration step using Polyethersulfone (PES) filters (PALL).

#### Purification of ConM and ConS Env trimers

Fed-batch supernatants were concentrated (≤ 14-times) and dia-filtered (≥ 6-times buffer exchange) into 20 mM Tris / 500 mM NaCl / pH 8.3 using Sartocon ECO (30 kDa MWCO, 0.70 m²) PES membranes (Sartorius). Buffer exchanged supernatants were filtered using 0.8 + 0.45 µm Sartopore 2 filters (Sartorius) particulate contaminant removal and further sterile filtered using 0.2 µm Sartopore 2 filters (Sartorius), then divided into four parts for further processing. Virus inactivation was performed by addition of 1% (w/w) Triton X-100 and incubation at RT for 60-120 mins and solutions sterile filtered using 0.45 µm Sartopore 2 filters (Sartorius). Triton X-100 inactivated solutions were batch-wise loaded onto a MAb PGT145 immunoaffinity column for capture of ConM and ConS Env trimers. The affinity columns were prepared from an in-house prepared GMP stock of MAb PGT145 by coupling of the antibody to Toyopearl 650M chromatography resin (Tosoh). Loaded columns were washed with ≥ 10 GV of 20 mM Tris / 500 mM NaCl / pH 8.3 and bound ConM and ConS trimers were eluted from the columns with 50 mM histidine / 2 M MgCl_2_ / pH 6.0. Trimer elution fractions were concentrated (≤ 5-times) and dia-filtered (≥ 10-times buffer exchange) to 0.1 M Glycine / 10 mM Tris / pH 7.5 using a Sartocon Slice ECO (30 kDa MWCO, 0.14 m²) PES membrane (Sartorius). Trimer-containing fractions were pooled and passed over a pre-equilibrated MAbSelect SuRe Protein A (ProA) affinity column (GE Healthcare) to remove leached residual PGT145-affinity ligand from the previous column. The trimer containing ProA flow through fractions were loaded onto a Q-Sepharose FF anion-exchange column (GE Healthcare), and columns washed with ≥ 5 column volumes of 0.1 M Glycine / 0.01 M Tris / pH 7.5. Trimers were then actively eluted with 0.1 M Glycine / 0.01 M Tris / 0.25 M NaCl / pH 7.5 (injection grade). AEX eluate fractions were sterile filtered using 0.45 + 0.2 µm Sartobran 300 filters (Sartorius), concentrated and dia-filtered to 20 mM Tris / 150 mM NaCl / pH 7.5 (injection grade drug substance/product buffer) using a Sartocon Slice ECO (30 kDa MWCO, 0.14 m²), diluted to a final Env trimer concentration of 2 mg/mL and filtered using 0.45 + 0.2 µm Sartobran 300 filter capsules (Sartorius). As final steps, nano-filtration of formulated ConM and ConS drug substance batches was performed using Planova™ 20N (0.3 m²) (Asahi Kasei) filters for virus removal, and drug substance batches were further diluted to concentrations of ∼1 mg/mL in drug substance/drug product buffer (20 mM Tris / 150 mM NaCl / pH 7.5) and stored for further processing at 2-8°C.

#### EDC cross-linking of GMP produced trimers and PGT145 immunoaffinity purification

ConM and ConS trimer preparations manufactured under GMP conditions were cross-linked by EDC / sulfo-NHS chemistry. As a preparatory step, ConM and ConS bulk drug substance (1 mg x mL^-1^; 20 mM Tris / 150 mM NaCl / pH 7.5; injection grade) were concentrated and dia-filtered into their optimal cross-linking buffer environments. ConM bulk drug substance (DS) preparations were buffer exchanged to 20 mM HEPES / 150 mM NaCl / pH 7.5; injection grade, and ConS DS preparations were buffer exchanged to phosphate buffered saline (PBS), pH 7.4; injection grade (≥ 10 volume changes) using a Sartocon Slice ECO (30 kDa MWCO, 0.14 m²) PES membrane (Sartorius). Buffer exchanged DS preparations were further adjusted to a concentration of ∼1 mg x mL^-1^ in 20 mM Tris / 150 mM NaCl / pH 7.5; injection grade. EDC cross-linking conditions were different for ConM and ConS trimers. For both trimers, EDC and NHS stock solutions were diluted in the trimer specific buffers and mixed with equal amounts (w/w) of buffer exchanged trimer solution (1 mg x mL^-1^). Cross-linking reactions for ConM were performed at RT for 30 mins, and ConS was incubated for 10 mins. Optimal molar concentrations for cross-linking of ConM and ConS were 1 M EDC / 10 mM NHS and 0.25 M EDC / 10 mM NHS respectively. Cross-linking reactions were quenched by addition of equal amounts (w/w) of a 1 M Glycine, pH 7.4 stock solution and incubation at RT for ≥ 10 mins. Further, cross-linked ConM (ConM-EDC) and ConS (ConS-EDC) preparations were concentrated and dia-filtered (≥ 7-times buffer exchange) with 20 mM Tris / 500 mM NaCl / pH 8.3; injection grade and passed through a 0.45 + 0.2 µm Sartopore 2 300 (Sartorius) filter for sterile filtration. ConM-EDC and ConS-EDC trimer preparations were loaded onto a GMP grade MAb PGT145 immunoaffinity column for positive selection of well-folded / cross-linked ConM and ConS Env trimers as previously described for purification of non-cross-linked trimer preparations. Immediately after, ConM-EDC and ConS-EDC MAb PGT145 immunoaffinity eluates were concentrated (≤ 5-times) and dia-filtered (≥ 14-times buffer exchange) into drug substance (DS) formulation buffer 20 mM Tris / 150 mM NaCl / pH 7.5; injection grade. As a last step, formulated ConM-EDC and ConS-EDC solutions were 0.2 µm filtered using a PES beaker filter (Millipore), concentrations were adjusted to ∼1 mg x mL^-1^ and the final DS preparation were stored at 2-8 °C.

#### PGT145 ELISA for quantification of ConM and ConS from supernatant samples

96-well plates (Nunc, Maxisorp) were coated with 1 µg x mL^-1^ of MAb 2G12 (Polymun Scientific) in carbonate/bicarbonate coating buffer, pH 9.5 at 4°C overnight. Before use, plates were blocked with PBS + 0.1% Tween and 1% BSA (PBS-T, 1% BSA) at RT for 1h. Supernatant samples and reference standards were diluted performing a 1:2 serial dilution series in PBS-T, 1% BSA. Assay plates (pre-coated) were washed (4-times) with PBS-T and aliquots of serially-diluted samples and standards were transferred to pre-coated assay plates and incubated at RT for 1h. After washing (4-times with PBS-T), biotinylated MAb PGT145 at 1 µg/mL was applied for 1h. Following incubation with MAb PGT145, plates were washed and incubated with streptavidin-HRP conjugate (Roche) for 30 mins. Color development was started by addition of o-phenylenediamine dihydrochloride (OPD) substrate, the reaction stopped by addition of 25% H_2_SO_4_ (Merck) and OD492 was measured with a Synergy 2 plate reader (BioTek). Evaluation of unknown supernatant samples by interpolation from standard curves of ConM and ConS reference material was performed using the Gen5 Microplate Reader and Imager Software package, version 3.05 (BioTek).

#### Characterization of GMP-produced ConM, ConM-EDC, ConS and ConS-EDC trimer preparations

Trimer integrity and stability of GMP produced ConM and ConS trimer preparations and their EDC cross-linked versions were subject to long-term stability monitoring of up to 60 months. Trimer preparations were evaluated by BN PAGE (Native PAGE Novex 3–12% Bis-Tris Gels) and Size Exclusion HPLC (SE-HPLC) using 2 consecutive Acquity UPLC protein columns (Waters) with 200 Å and 450 Å to asses integrity and purity of GMP trimer preparations. Reducing and non-reducing SDS PAGE (NuPAGE 4 to 12%, Bis-Tris gels) was performed together with immunoblotting and probing with anti-Env specific antibodies (5F3, 447-52D, 2G12). GMP produced preparations were routinely monitored for the presence of particulate contaminants, pH, endotoxin levels and osmolality.

#### DSC analysis

Thermal denaturation of unmodified and cross-linked GMP batches of ConM and ConS was studied using a nano-DSC calorimeter (TA instruments, Etten-Leur, The Netherlands)[39]. Briefly, trimers were first dialyzed against PBS and concentration adjusted to ∼0.25 or ∼1.0 mg/mL, respectively. After sample loading, thermal denaturation was probed at a scan rate of 60 °C/h. Buffer correction, normalization, and baseline subtraction procedures were applied and data were analyzed using the NanoAnalyze Software v.3.3.0 (TA Instruments). The data were fitted using a non-two-state model, as the asymmetry of some of the peaks suggested that unfolding intermediates were present. DSC experiments were performed with a D7324-tagged trimer, but the presence of the D7324-tag did not alter the *T*_m_ values compared to the corresponding non-tagged trimers [39].

#### Free amine assay

Unmodified or cross-linked trimer (5 μg) in 20 μL PBS was added to 30 μL of 0.1 M NaHCO_3,_ pH 8.5. 25 μL of 5% 2,4,6-Trinitrobenzene Sulfonic Acid (TNBSA) diluted 1/500 in 0.1 M NaHCO_3_ pH 8.5 was added to the samples for 2 h at 37°C, followed by 25 μL of 10% SDS and 12.5 μL of 1M HCl. Samples were vortexed and the optical density read at 335 nm. The relative quantity of free amines was calculated as (OD_335_ (GLA-SOSIP trimer)– OD_335_ (blank)) / (OD_335_ (SOSIP trimer)–OD_335_ (blank)).

#### Capture ELISA for Env trimer binding by human mAbs

ELISA plates (Greiner Bio-One) were coated with 4 μg/mL of capture mAb 2G12 at 4°C overnight in PBS. After blocking with 2% BSA/PBS + 0.05% Tween, trimer (0.2 μg/mL) were captured, labelled with a titration series of biotinylated human mAbs followed by peroxidase-conjugated streptavidin detection reagent (Jackson ImmunoResearch). The colorimetric endpoint was obtained using the one-step ultra TMB substrate (Thermofisher). MAbs were developed until a signal of approximately 1–2 optical density (OD_450_) units was generated for each antibody, leading to longer incubation periods for non-Nabs, and colour development was stopped with sulfuric acid (0.5 M) and the OD_450-570_ measured. All ELISA signals were corrected by subtracting the background signal obtained in the absence of primary antibody and the resulting data were plotted against the log_10_ of the antibody concentration using GraphPad Prism V7.0. To generate binding indices from ELISA titration curves, an area under the curve (AUC) analysis of ligand-trimer binding was performed; the binding index represents the ratio of cross-linked trimer value to the value of the matched unmodified SOSIP trimer that was used for cross-linking. Binding indices were calculated as (AUC(GLA-SOSIP trimer)— AUC(blank)) / (AUC(SOSIP trimer)—AUC(blank)), where blank = negative control curve of the respective mAb without antigen. Indices <1 indicate reduced binding to the cross-linked trimer compared to its unmodified counterpart, and the converse for values >1.

#### Env trimer ELISA for mouse sera

Detection of mouse trimer-specific serum antibodies was performed using endpoint titer ELISA. 2G12 antibody (4 μg/mL, 50 μL/well) was captured overnight at 4°C onto high-protein-binding ELISA plates (Spectraplate 96HB, Perkin Elmer). Plates were washed in PBS/Tween (0.05% v/v) and wells blocked using BSA (2% w/v; 200 μL/well) for 1 h at RT, and washed. Unmodified trimer (0.2 μg/mL, 50 μL/well) was added for 2 h at RT and washed as before. Mouse serum samples diluted in PBS/BSA (1% w/v) starting at 1:100 then stepwise 5-fold were added to the ELISA plates (50 μL/well) and incubated overnight at 4°C. Plates were washed and Peroxidase-conjugated rabbit anti-mouse IgG antibody (1:5000; 50 μL/well, Jackson Immunoresearch) added to all wells for 1 hr at RT. Plates were washed and TMB substrate (50 μL/well, Thermofisher Scientific) added to all wells. Colour development was monitored and terminated after 10 mins using sulphuric acid (0.5 M; 50 μL/well). Optical density (OD) values for each well were calculated as OD_450-570nm_. Background values (OD_no serum_) were subtracted from sample readings. Endpoint titers were calculated using non-linear regression curve fitting and interpolated values transformed into log_10_ endpoint titers (Graphpad Prism 7 for Mac).

#### Biolayer interferometry (BLI) trimer antigenicity analysis

The GMP trimer constructs in this study are devoid of tags, therefore Fab fragments were loaded onto anti-Human Fab CHI (FAB2G) biosensors and dipped into varying concentrations (in nM: 2000, 1000, 500, 250, 125, 62.5, 31.25) of trimer using an OctetRed 96 instrument (ForteBio). Trimers and Fabs were diluted into 1X kinetics buffer (PBS pH 7.4 + 0.01% BSA, 0.002% Tween-20), and allowed to reach room temperature before beginning the assay. Association was measured for 600 seconds, followed by dissociation for 1200 seconds in 1X kinetics buffer. A reference well containing kinetics buffer was subtracted from each dataset, curves were aligned on the y axis using the baseline step, and an inter-step correction was applied between the association and the dissociation curves. A 1:1 binding model, which assumes first-order kinetics and each binding site on the SOSIP binds to immobilized Fab at an equal rate, was fitted to the data and used to determine kinetic parameters.

#### Analysis of elastase and CathepsinG activity on trimers

To test Env stability in the presence of neutrophil-released proteases, 0.01 Units of elastase or cathepsinG (Sigma) was added to 1 μg of each trimer in a final volume of 11.1 μL PBS and incubated at 37° C for 1 or 24 h. A negative control, containing no enzyme, also underwent a 24 h incubation period at 37° C. After protease exposure the samples were analysed by reducing and denaturing polyacrylamide gel electrophoresis on 4-12% bis-tris gels (Life Technologies). Protein bands were developed with Simply Blue Safe Stain (Life Technologies) and band intensity analysed on Biorad GelDoc XR with Lab Image software. Band intensity of the untreated control was set at 100% and all other samples were normalised to that.

#### TZM-bl neutralization assay

Infectious molecular clones (IMC) ConM and ConS [4] were produced in HEK293T cells, titered and used in TZM-bl assay to determine nAb responses as previously described[55]. Briefly, duplicates of six steps of 3-fold dilution, starting with 1:20 of each serum, were incubated with viral supernatant (at relative luminescence units (RLU) between 150.000 and 200.000) for 1 h. Thereafter, 10^4^ TZM-bl cells were added, and plates incubated for 48 h at 37°C, after which Bright-Glo Luciferase assay system (Promega, Madison, Wisconsin, USA) was added to measure luciferase activity with a Mithras luminometer (Berthold, Germany). Positive controls were sera of HIV-1-infected individuals and monoclonal antibodies with known neutralizing titers. Neutralization titers were defined as the sample dilution at which RLU were reduced by 50% compared to virus control wells after subtraction of background RLU in control wells with only cells. Inhibitory concentrations (IC) 50 were calculated with a linear interpolation method using the mean of the duplicate responses [55].

#### T cell cytokine responses

Spleens were harvested from mice 4 weeks after the final boost, dissected using aseptic technique, and single cells isolated by passing through a 100 μm filter. Splenocytes were resuspended in supplemented RPMI (10% fetal bovine serum, 10 mM HEPES, 2 mM Glutamax, 1x Penicillin-Streptomycin, 50 μM 2-ME) and plated at 500,000 splenocytes per well in 96 U-bottomed plates in a final volume of 200 μL. Splenocytes were either pulsed with relevant antigen at 50 μg/mL or no antigen as a control. On day 3, 100 μL supernatant was harvested and frozen at −80 °C for cytokine analysis. The concentration of IL-2 in 2x diluted supernatants was measured using an IL-2 mouse ELISA kit (Thermo Fisher Scientific) as per the manufacturer’s instructions. The concentration of IL-2, IFNγ, IL-5, IL-13 and IL-17a in supernatants was measured using a Luminex multiplex assay (R&D) performed according to the manufacturer’s instructions. Briefly, supernatant was diluted 1:2 in RD1W buffer, whilst the standards were prepared as instructed. 50 μL of standard or diluted supernatant was pipetted into the wells of a black 96-well plate and 50 μL of magnetic-bead cocktail was added to each well. The plates were then incubated for 2 h on a shaker set to 800 rpm. The supernatant was removed and wells washed 3x with 300 μL of wash buffer using a magnetic plate washer. 50 μL of biotin-labelled secondary antibody, diluted as per the instructions was added to the wells and the plate returned to the shaker for 1 h. The plates were washed again before a 30 min incubation with streptavidin-PE, washed and the plates read on a Luminex Bio-Plex (Bio-Rad).

#### Negative Stain EM

GMP SOSIPs were diluted in TBS pH 7.4 to ∼0.02 mg/mL, applied to glow-discharged copper EM grids containing a continuous carbon film, then negatively stained with 2% uranyl formate for ∼10 seconds. Micrographs were collected on a Talos 200C transmission electron microscope (ThermoScientific) operated at 200 keV, at a nominal magnification of 73,000X resulting in a pixel size of 1.98Å. CTF estimation was performed with CTFFIND4 [56], and then all subsequent processing steps were carried out in RELION-3 [57]. Particles were picked with the RELION Gaussian picker, and an initial 2D classification was performed to eliminate noise and non-protein particle picks. A second round of classification was then used to determine the percent of trimeric and properly folded SOSIPs within each GMP sample. The percent well-folded trimers was reported as the number of particles contained within 2D classes of trimeric SOSIPs, divided by the total number of particles in the particle stack.

#### Cryo-electron microscopy sample preparation

Unmodified and cross-linked GMP SOSIP trimers were incubated with a 6-fold molar excess of PGT122 Fab overnight at 4°C. Excess Fab was removed by ultrafiltration in TBS pH 7.4 with a 100 kDa cutoff Centricon filter (Amicon Ultra, Millipore), and the sample was concentrated to ∼5 mg/mL for cryo grid freezing. Immediately before vitrification, *n*-Dodecyl β-D-maltoside (DDM) was added to a final concentration of 0.25 mM, which improved the angular distribution of the protein particles in vitreous ice. 3 µL of this mixture was applied to Quantifoil R1.2/1.3 holey carbon Cu 400 mesh grids, and blotted and plunge-frozen in liquid ethane with the Vitrobot Mark IV.

#### Cryo-electron microscopy data collection

Micrographs were collected on a Titan Krios (ThermoScientific) operating at 300 keV coupled with a Gatan K2 direct electron detector via the Leginon interface[58]. Each exposure image was collected in counting mode at 29000X nominal magnification resulting in a pixel size of 1.03 Å/pixel, using a dose rate of ∼5.5 e-/pix/sec, and 200 ms exposure per frame. The total dose on the sample for each movie micrograph was 50 e-/Å^2^, and the nominal defocus range used was −1.0 to −2.2 μm. These imaging conditions were kept consistent for each of the four samples (ConS, ConS-EDC, ConM, and ConM-EDC).

#### Cryo-electron microscopy data processing

Movie micrograph frames were aligned and dose-weighted using MotionCorr2 [59], and imported into cryoSPARC v2 [60]. CTF models were calculated using CTFFIND4. The cryoSPARC blob picker was used on a subset of micrographs to generate an initial set of particle picks, which were subjected to 3 rounds of 2D classification. High quality and representative classes were selected and used as 2D templates for template picking on the entire dataset. The resulting particle images were subjected to multiple rounds of 2D classification, followed by one round of *ab initio* reconstruction to generate an initial model. This map was used as an initial model for one round of heterogeneous refinement, specifying two or four classes. For each data set there was no observable heterogeneity, but this step served to further eliminate low resolution particles. Particles from the dominant class(es) were re-extracted with a box size of 352 pixels, and subjected to homogeneous refinement. Next, iterative rounds of global CTF refinement followed by homogenous refinement were performed until the resolution had converged, followed by a final round of non-uniform refinement. The ConS data showed potential to reach higher resolution, therefore these data were first processed in RELION according to standard single particle workflows in order to perform Bayesian polishing. This map reached a resolution of 3.3Å, but PGT122 Fab density was poorly resolved. Polished particles were extracted and imported into cryoSPARC v2, and the same workflow as described above was performed for this particle stack. This procedure led to a very high-quality reconstruction throughout the complex, with a final resolution of 3.1Å. For unmodified ConM, higher resolution was achieved by importing frames into cryoSPARC2 and using the native patch motion correction utility. This allowed for local (single particle) motion to be corrected, which improved the resolution and led to the final 3.4Å reconstruction. For EDC cross-linked ConS and ConM, these procedures led to no further improvements, thus for these data pre-aligned and dose-weighted micrographs were imported, as described above. All resolutions are reported according to the “gold standard” 0.143 FSC cut-off [61].

#### Model building

The initial model for ConM-PGT122 comprised the ConM SOSIP coordinates from the ConM-PGT122-35022 crystal structure (PDB 6IEQ), and the PGT122 Fab coordinates from a crystal structure of BG505-PGT122-35022 (PDB 4TVP). These coordinates served as the template for complete rebuilding and all-atom refinement with RosettaCM [62], which couples the fragment-based *de novo* Rosetta builder with sequence and structural constraints from the template(s), as well as a user-defined electron density weight. The lowest energy model was selected and glycans were added to the model in Coot [63], which was then iteratively refined in real-space with Coot and PHENIX [64]. Once the model had converged, disordered regions were removed and a final high resolution, all-atom refinement with Rosetta Relax [65] was performed. This ConM-PGT122 cryo-EM structure then served as the initial model for both ConM-EDC and unmodified ConS, which followed a nearly identical procedure: full model rebuilding with RosettaCM, real-space refinement in Coot and PHENIX, and final refinement with Rosetta Relax. This same procedure was also used for the ConS-EDC structure, except that the unmodified ConS structure was used as the initial model. Structure validation was performed with MolProbity [66], PHENIX, and EMRinger [67].

#### RMSD analysis

Alpha carbon (Cα) root-mean-square deviation (RMSD) was calculated by superimposition of structures in UCSF Chimera [68]. This was performed globally and locally, in order to differentiate global conformational changes from local changes in structure of specific epitopes. Values for global RMSD are listed in Figures 2B-E, and Supplementary Figure 7. The local Cα RMSD was mapped onto the cryo-EM structures in Figure 2B-E using UCSF Chimera.

#### Mouse immunizations

All experiments used 8–12 week old female BALB/c mice (Charles River) maintained under specific pathogen-free conditions in the Sir William Dunn school facility. In two independent experiments 4 or 5 mice per group were immunized by subcutaneous administration with 10 μg trimer formulated in 100 μL PBS/MPLA (10 µg/dose) formulation on weeks 0, 4 and 8. Mice were monitored for adverse symptoms throughout. Blood was collected by tail bleed on weeks −1, 2, 6, 10 and 12, serum separated and stored at −20°C.

#### Rabbit immunizations, IgG purification and ELISA

A total of 30 New Zealand White Specific Pathogen Free (SPF) rabbits (23 males and 23 females (were nulliparous and non-pregnant)), 11 weeks old and weighing approximately 2 kg, were immunised intramuscularly at the Research Toxicology Centre S.p.A, Italy. Each study group (detailed in Supplementary Table 3) consisted of 3 male and 3 female animals. The rabbits received a total of 4 administrations (Weeks 0, 3, 6 and 9) into the lateral surface of the quadriceps muscle of both legs. Rabbit serum was isolated from blood at 0, 3, 6, 9- and 12-weeks first post-vaccination, heat inactivated, aliquoted and stored at −20°C prior to assessment for antigen-specific IgG. Antigen-specific gp140 binding antibodies were measured using standardised ELISA platforms. In serum samples, antigen-specific IgG was measured. In brief, 96-well medium binding plates (Griener, Kremsmunster, Austria) were coated with either recombinant ConM, ConM-edc, ConS or ConS-edc gp140 (Polymun Scientific) at 1 ug/mL in PBS for 1 h 37°C. As reference material, standard immunoglobulins (Serotec, UK) were captured with anti-Rabbit specific goat antibodies (Millipore, UK). After blocking with assay buffer (5% bovine serum albumin; Sigma–Aldrich; 0.05% Tween ThermoFisher Scientific, Pittsburgh, PA), samples were initially screened at 1:100 dilution (then titrated to optimal dilutions) on antigen-coated wells, and serial dilutions of immunoglobulin standards were added to the anti-Rabbit capture antibody coated wells and incubated for 1 h at 37°C. Secondary antibody HRP-conjugated anti-rabbit IgG was added at 1:20,000 dilution and incubated for 1 h at 37°C. Plates were developed with SureBlue TMB substrate (KPL, Insight Biotechnology, London, United Kingdom). The reaction was stopped after 5 min by adding TMB stop solution (KPL, Insight Biotechnology), and the absorbance was read at 450 nm on a VersaMax 96-well microplate reader (Molecular Devices, Sunnyvale, CA). The ELISA data are expressed as positive if the blank subtracted OD450 nm was above the predetermined cut-off of OD 0.2 nm and values are on the linear range of the curve. To ensure assay sensitivity, a positive control composed of gp140 positive pooled rabbit serum samples and a negative control composed of anti-Ad4 Rabbit Hypersera (PAXVAX, San Diego, US) were used. Analyses of the data were performed using SoftMax Pro GxP software v6.5 (Molecular Devices).

#### Ethics statement

Animal research using rabbits and mice was carried out in full accordance with local and national ethical guidelines. All protocols for breeding and procedures with mice were approved by the Home Office UK, under the Animals (Scientific Procedures) Act 1986 and Home Office license PPL3003421. Rabbit studies were carried out at Covance Inc.

#### Statistical analyses

Statistical analysis was performed in Prism using the tests described in the corresponding figure legends. Briefly, one-way ANOVA of log-transformed data with Sidak’s post-test correction to account for multiple comparisons were used to analyze normally-distributed data including log-transformed endpoint titers. Non-parametric analysis (not assuming a Gaussian distribution) between two independent groups was performed using a two-tailed Mann-Whitney U test and an unmatched, unpaired Kruskal-Wallis test with Dunn’s multiple comparison test was used to compare non-normally distributed data with more than one comparison.

## Acknowledgements

We thank John Mascola, Peter Kwong, Dennis Burton, Michael Nussenzweig, Mark Connors and James Robinson for donating antibodies and other reagents either directly or through the NIH AIDS Reagents Program. We thank Dennis Burton and the IAVI Neutralizing Antibody Consortium for reagents. QJS is a Jenner Institute Investigator and James Martin School Senior Fellow.

## Author contributions

Gregory Martin, Conceptualization of the work, Data collection, Data analysis and interpretation, Writing – original draft, writing-review and editing, Final approval of the version to be published.

Rebecca A Russell, Conceptualization of the work, Data collection, Data analysis and interpretation, Supervision, Writing – review and editing, Final approval of the version to be published.

Philip Mundsperger, Data collection, Data analysis and interpretation, Writing – review and editing, Final approval of the version to be published.

Scarlett Harris, Data collection, Data analysis and interpretation, Writing – review and editing, Final approval of the version to be published.

Lu Jovanoska, Data collection, Data analysis and interpretation, Writing – review and editing, Final approval of the version to be published.

Luiza Farache Trajano - data collection, data analysis and interpretation

Leon Macfarlane - Data collection, Data analysis and interpretation, Final approval of the version to be published.

Hannah Cheeseman, Data collection, Data analysis and interpretation, Writing – review and editing, Final approval of the version to be published.

Katalin Fabian, Data collection, Data analysis and interpretation, Writing – review and editing, Final approval of the version to be published.

Monica Tolazzi, Data collection, Data analysis and interpretation, Writing – review and editing, Final approval of the version to be published.

Marielle Breemen, Data collection, Data analysis and interpretation, Writing – review and editing, Final approval of the version to be published.

Torben Schiffner, Conceptualization of the work, Writing – review and editing, Final approval of the version to be published.

Kwinten Sliepen, Conceptualization of the work, Data collection, Data analysis and interpretation, Supervision, Writing – review and editing, Final approval of the version to be published.

Yoann Aldon, Conceptualization of the work, Writing – review and editing, Final approval of the version to be published.

Gabriella Scarlatti, Conceptualization of the work, Data analysis and interpretation, Supervision, Writing – review and editing, Final approval of the version to be published.

Rogier W Sanders, Conceptualization of the work, Funding acquisition, Writing – review and editing, Final approval of the version to be published.

Robin Shattock, Conceptualization of the work, Funding acquisition, Writing – review and editing, Final approval of the version to be published.

Dietmar Katinger, Funding acquisition, Project administration, Supervision, Writing – review & editing, Final approval of the version to be published.

Andrew B Ward, Funding acquisition, Project administration, Supervision, Writing – review & editing, Final approval of the version to be published.

Quentin J. Sattentau, Conceptualization, Formal analysis, Funding acquisition, Project administration, Supervision, Writing – original draft, Writing – review & editing, Final approval of the version to be published.

## Data availability

The final cryo-EM reconstructions and the resulting structural models have been deposited into the Protein Data Bank (PDB) the Electron Microscopy Data Bank (EMDB) under the following accession codes: **ConS:** PDB 7LX2, EMDB 23564; **ConS-EDC:** PDB 7LX3, EMDB 23565; **ConM:** PDB 7LXM, EMDB 23571; **ConM-EDC:** PDB 7LXN, EMDB 23572.

---

**Supplementary Figure 1.**
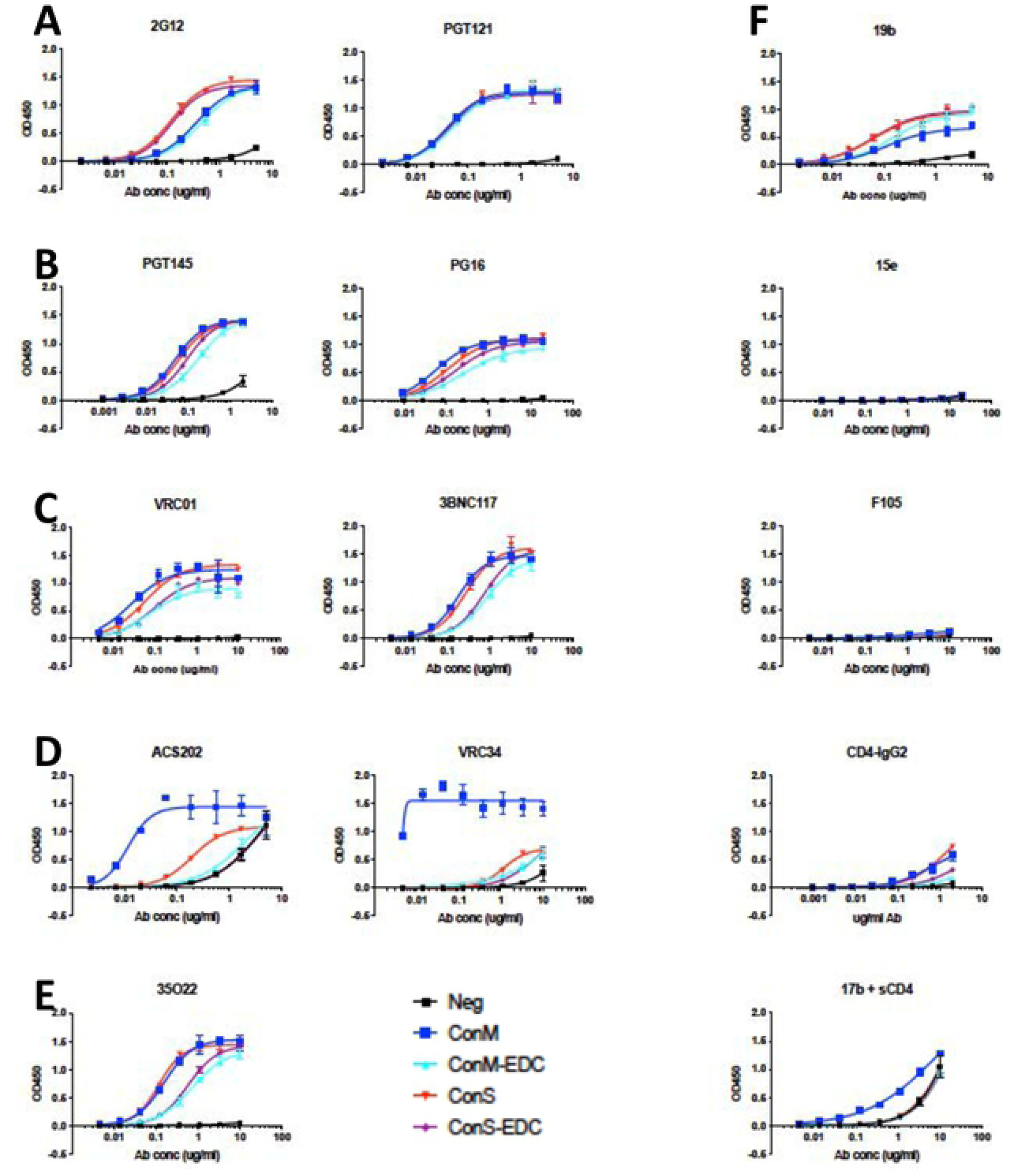
ELISA binding curves associated with Figure l F. **A**) V3 glycan bn Abs; **B**) Apex quaternary bnAbs; **C**) CD4bs bnAbs; **D**) fusion peptide bnAbs; **E**) gp120-gp41 interface bnAb; **F**) non-nAbs and CD4-lgG2.

**Supplementary figure 2.**
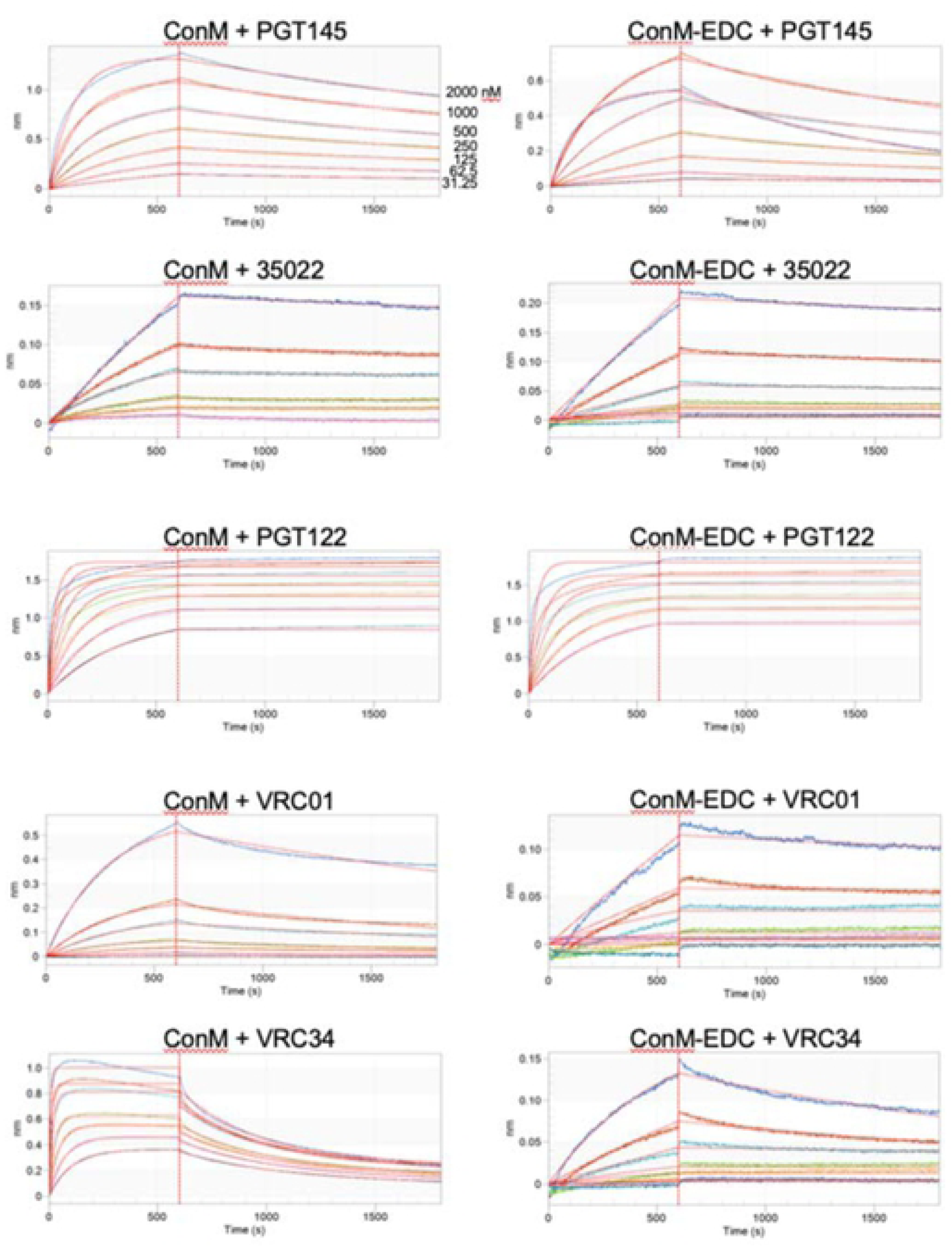
Impact of EDC on binding of bNAbs to ConM and ConM-EDC trimers assayed by biolayer interferometry, associated with >Table 1. The GMP trimers in this study are untagged, thus bnAb Fab fragments were immobilized to anti-human Fab-CHI biosensors, and binding was analyzed at a range of SOSIP concentrations (2000, 1000, 500, 250, 125, 62.5, 31.25 nM). The aligned and reference-subtracted binding response (colored traces) was fitted to a 1:1 binding model (red lines) to determine kinetic parameters. SOSIP concentrations which gave poor fits (R2<0. 95) were not used for the kinetic analysis. In each case the final values were the average of at least three trimer concentrations.

**Supplementary figure 3.**
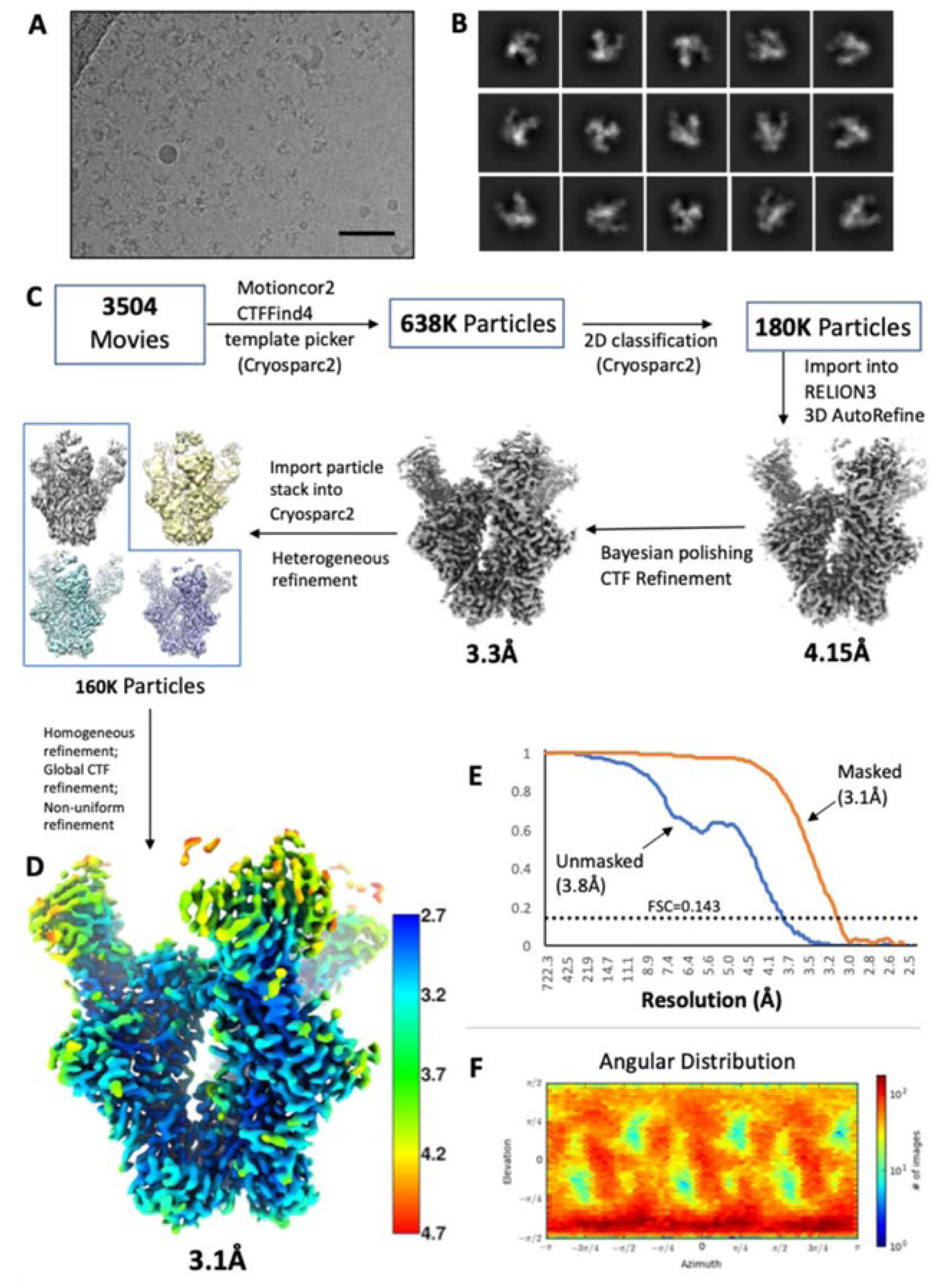
Cryo-EM data processing workflow and reconstruction for unmodified Cons. (**A**) Motion-corrected micrograph from Titan Krios (1.03 A/pixel). (**B**) Select 20 classes from Cryosparc v2. (**C**) Data processing workflow. (**D**) Final reconstruction and local resolution, calculated in Cryosparc v2. (**E**) Fourier shell correlation. Reported resolutions are according the 0.143 FSC gold standard. (**F**) Angular distribution plot of final reconstruction, from Cryosparc v2.

**Supplementary figure 4.**
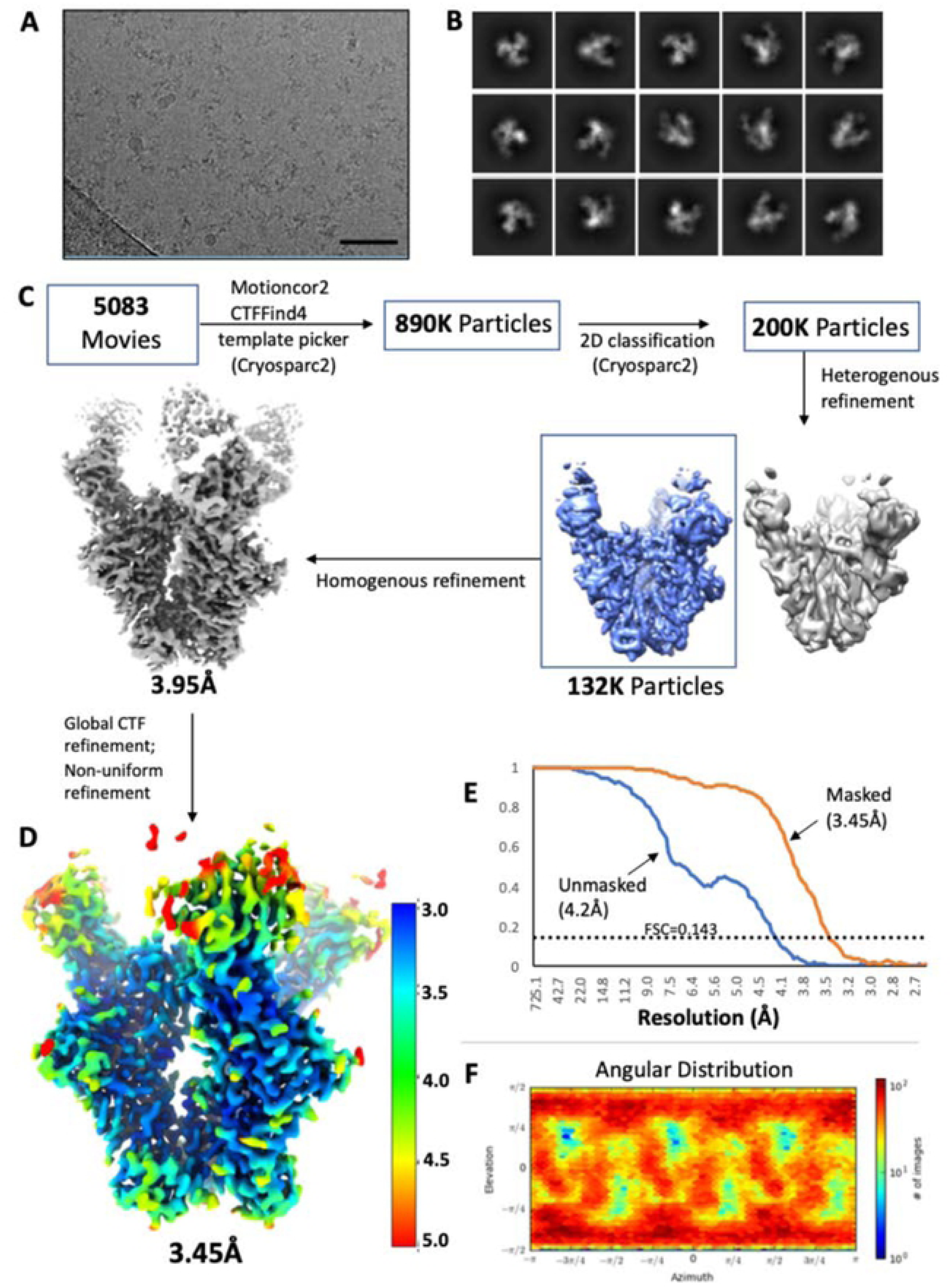
Cryo-EM data processing workflow and reconstruction for EDC-crosslinked Cons. (**A**) Motion-corrected micrograph from Titan Krios (1.03 A/pixel). (**B**) Select 20 classes from Cryosparc v2. (**C**) Data processing workflow. (**D**) Final reconstruction and local resolution, calculated in Cryosparc v2. (**E**) Fourier shell correlation. Reported resolutions are according the 0.143 FSC gold standard. (**F**) Angular distribution plot of final reconstruction, from Cryosparc v2.

**Supplementary figure 5.**
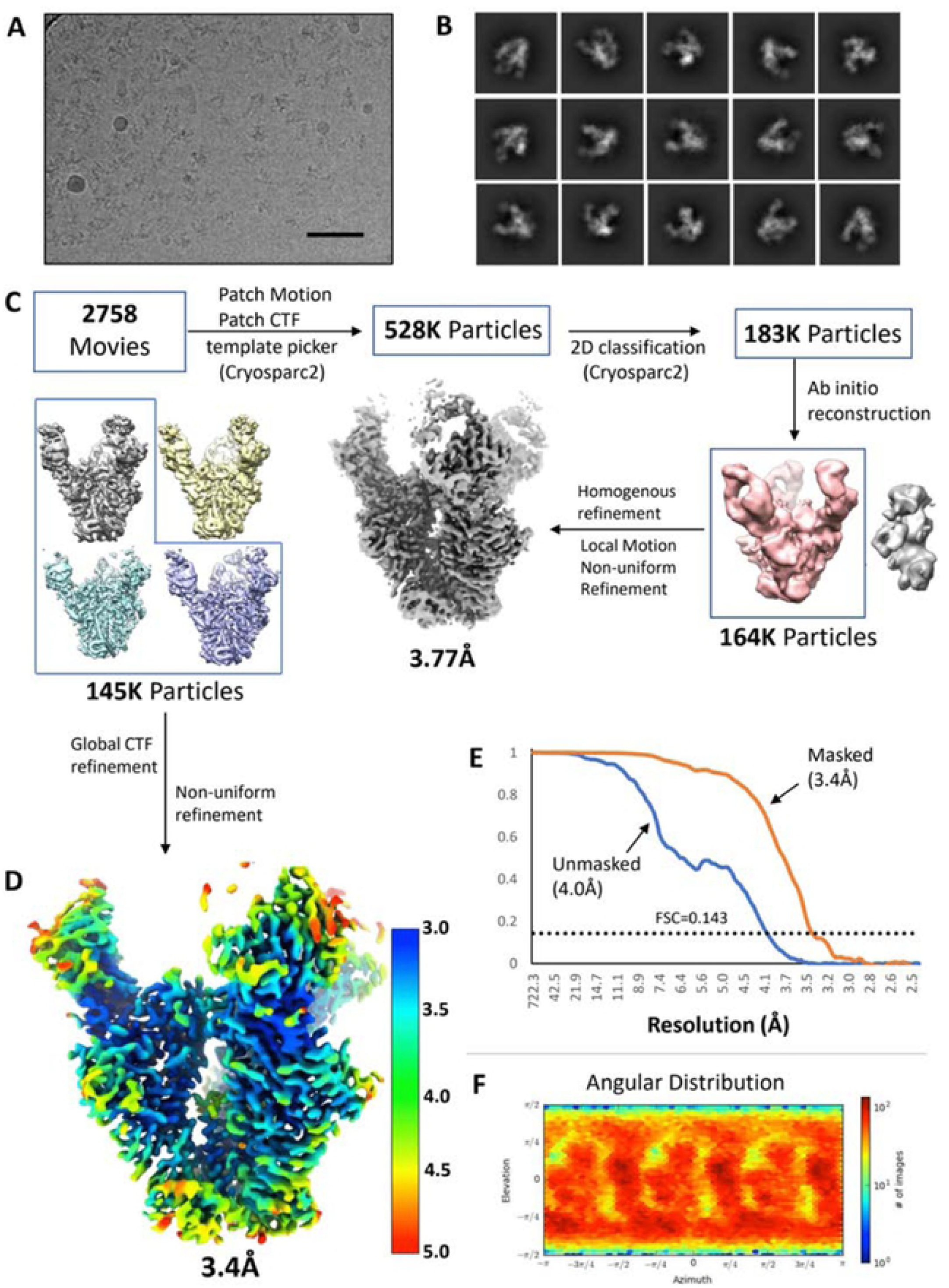
Cryo-EM data processing workflow and reconstruction for ConM. (**A**) Motion-correctedmicrograph from Titan Krios (1.03 A/pixel). (**B**) Select 20 classes from Cryosparc v2. (**C**) Data processing workflow. (**D**) Final reconstruction and local resolution, calculated in Cryosparc v2. (**E**) Fourier shell correlation. Reported resolutions are according the 0.143 FSC gold standard. (**F**) Angular distribution plot of final reconstruction, from Cryosparc v2.

**Supplementary figure 6.**
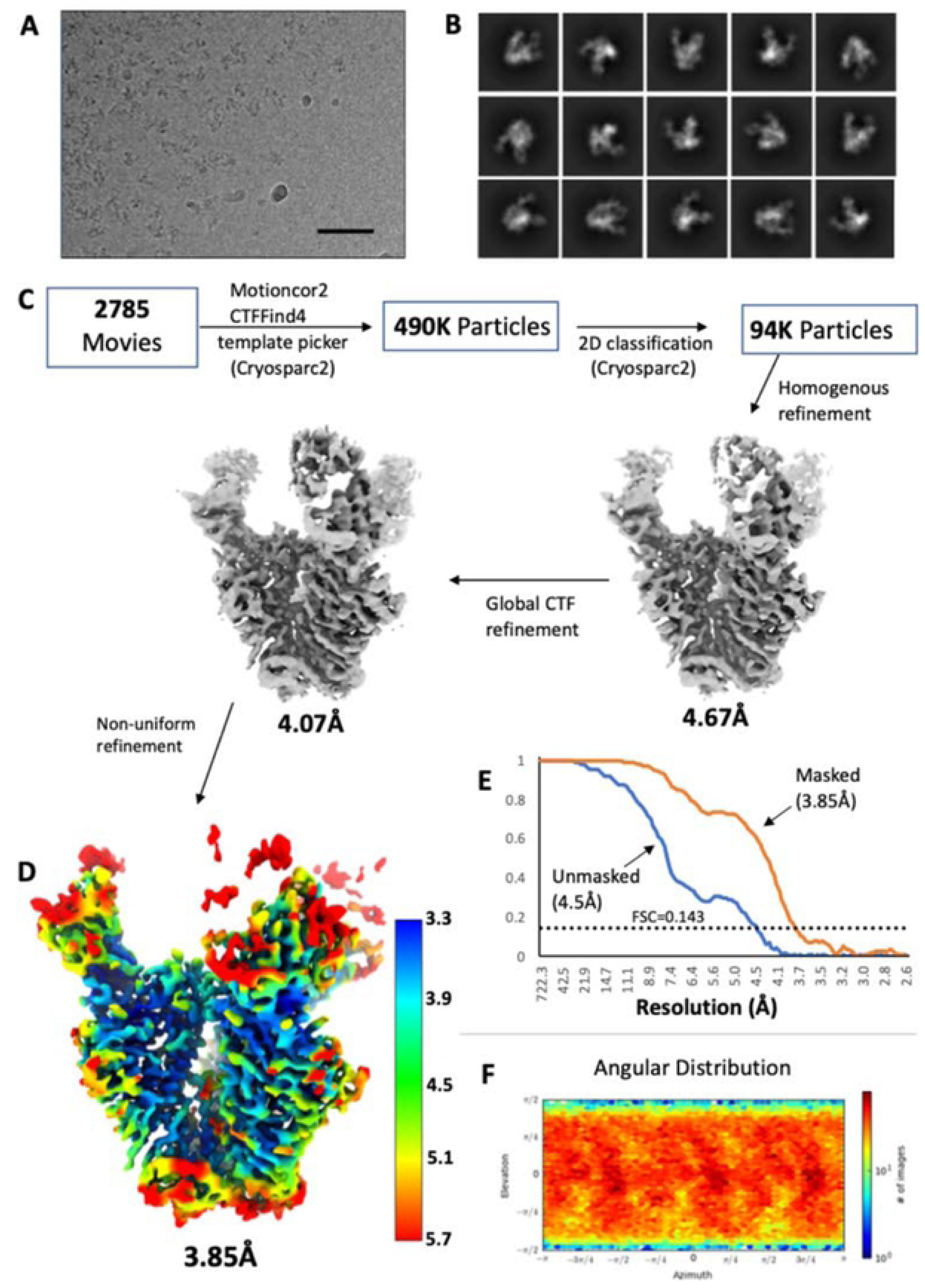
Cryo-EM data processing workflow and reconstruction for EDC cross-linked ConM. (**A**) Motion-corrected micrograph from Titan Krios (1.03 A/pixel). (**B**) Select 20 classes from Cryosparc v2. (**C**) Data processing workflow. (**D**) Final reconstruction and local resolution, calculated in Cryosparc v2. (**E**) Fourier shell correlation. Reported resolutions are according the 0.143 FSC gold standard. (F) Angular distribution plot of final reconstruction, from Cryosparc v2.

**Supplementary Figure 7.**
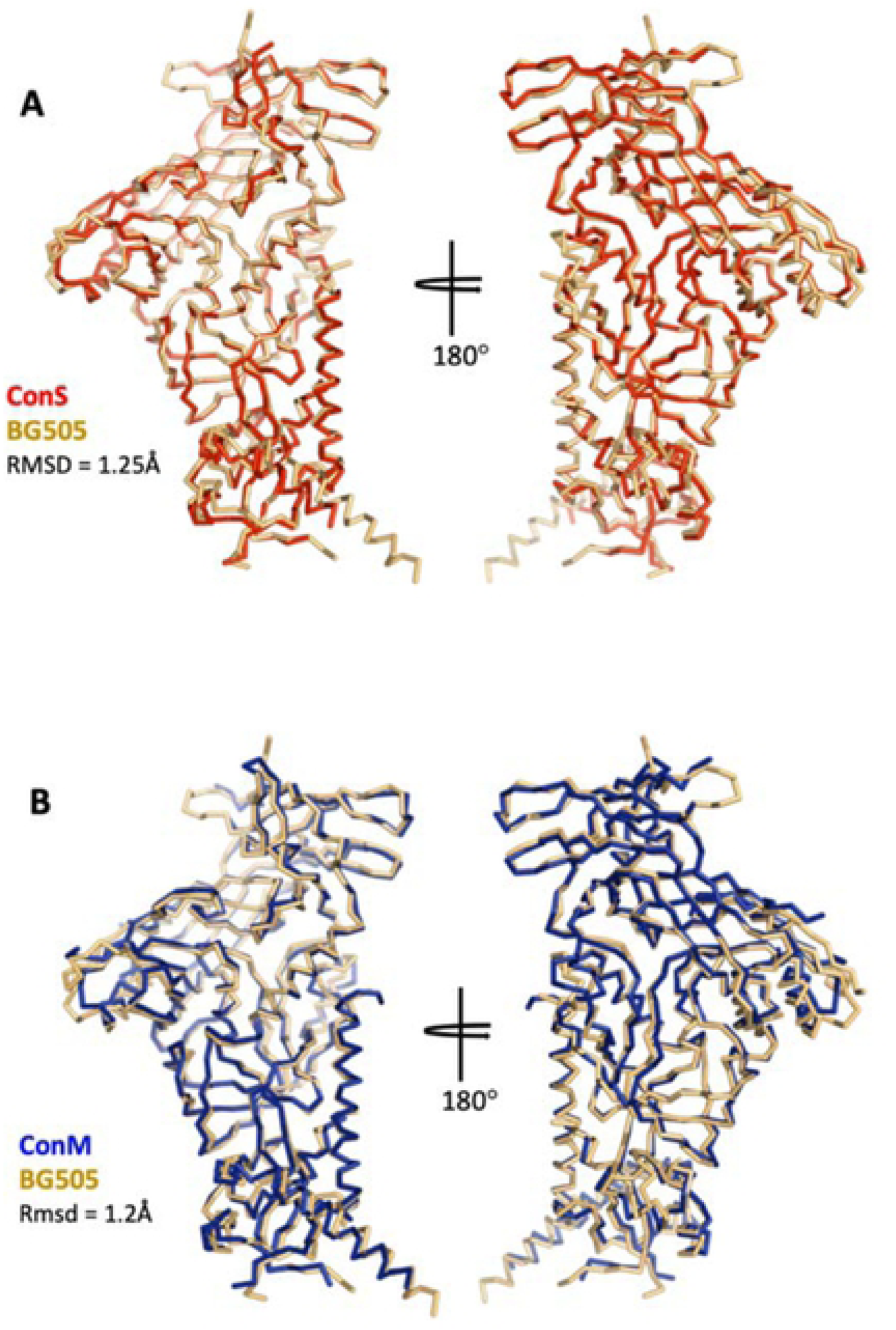
Structural comparisons of ConM and Cons with BG505. (**A**) Superimposition of a single protomer of the ConS-PGT122 cryo-EM structure with the BG505-PGT122-35022 X-ray structure (PDB 4TVP). Shown is a ribbon representation of the Ca chain trace. The Ca RMSD is 1.25A. (**B**) Same as in (**A**) but comparing BG505 with the ConM­ PGT122 cryo-EM structure. The Ca. RMSD is 1.2A.

**Supplementary Figure 8.**
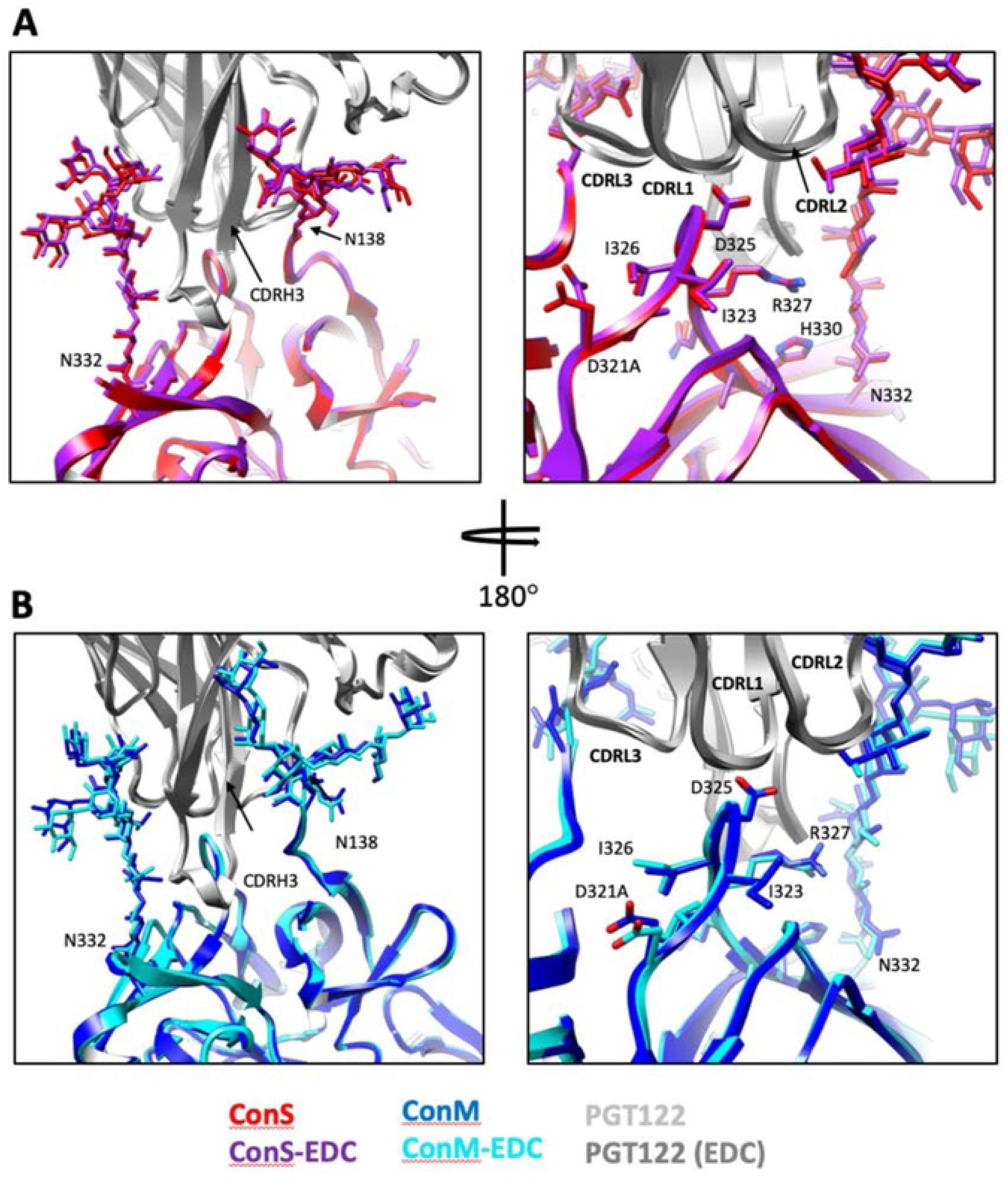
Impact of cross-linking on the PGT122 epitope. (**A**) Superimposition of Cons (red) and ConS-EDC (purple), showing zoomed-in view of PGT122 (grey) binding site. Glycans at N332 and N138 are also shown, which were highly-ordered in the cryo-EM structure. At right the view is rotated 180° and side chains of residues directly in contact with antibody are shown as sticks, including the G324DIR327 motif. (**B**) Same as in (**A**), but for ConM and ConM-EDC.

**Supplementary Figure 9.**
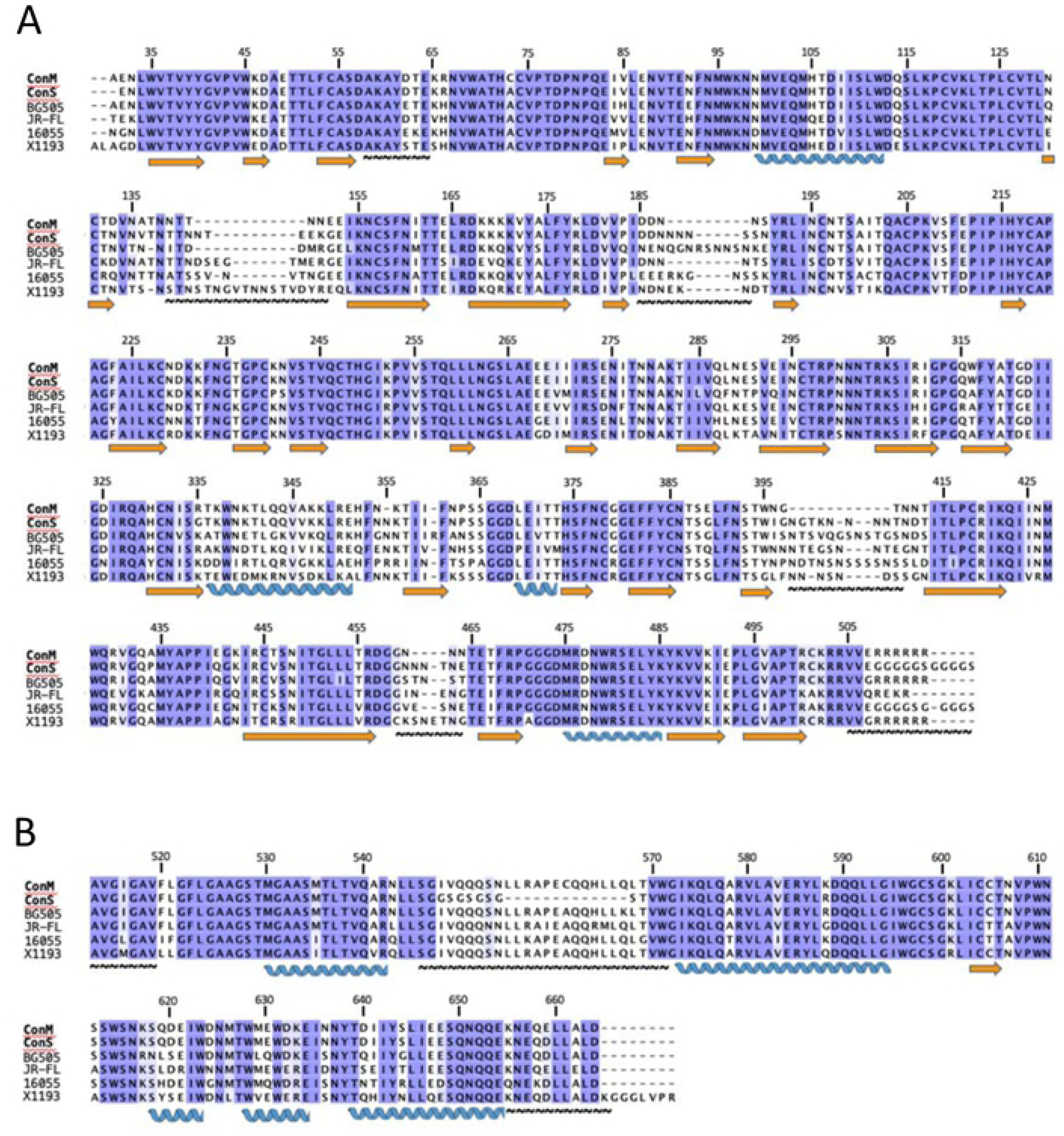
Sequence alignment of ConM and Cons with Env sequences from various clades. BG505, clade A; JR-FL, clade B; 16055 NFL TD, clade C; X1193.c1 SOSIP.664, clade G. (**A**) gp120 alignment (**B**) gp41 alignment. Orange arrows denote beta strands, blue coils denote alpha helices, and tilde stringsare regions mostly disordered in the ConM and Cons structures. Numberingis accordni g to the HXB2 system.

**Supplementary Figure 10.**
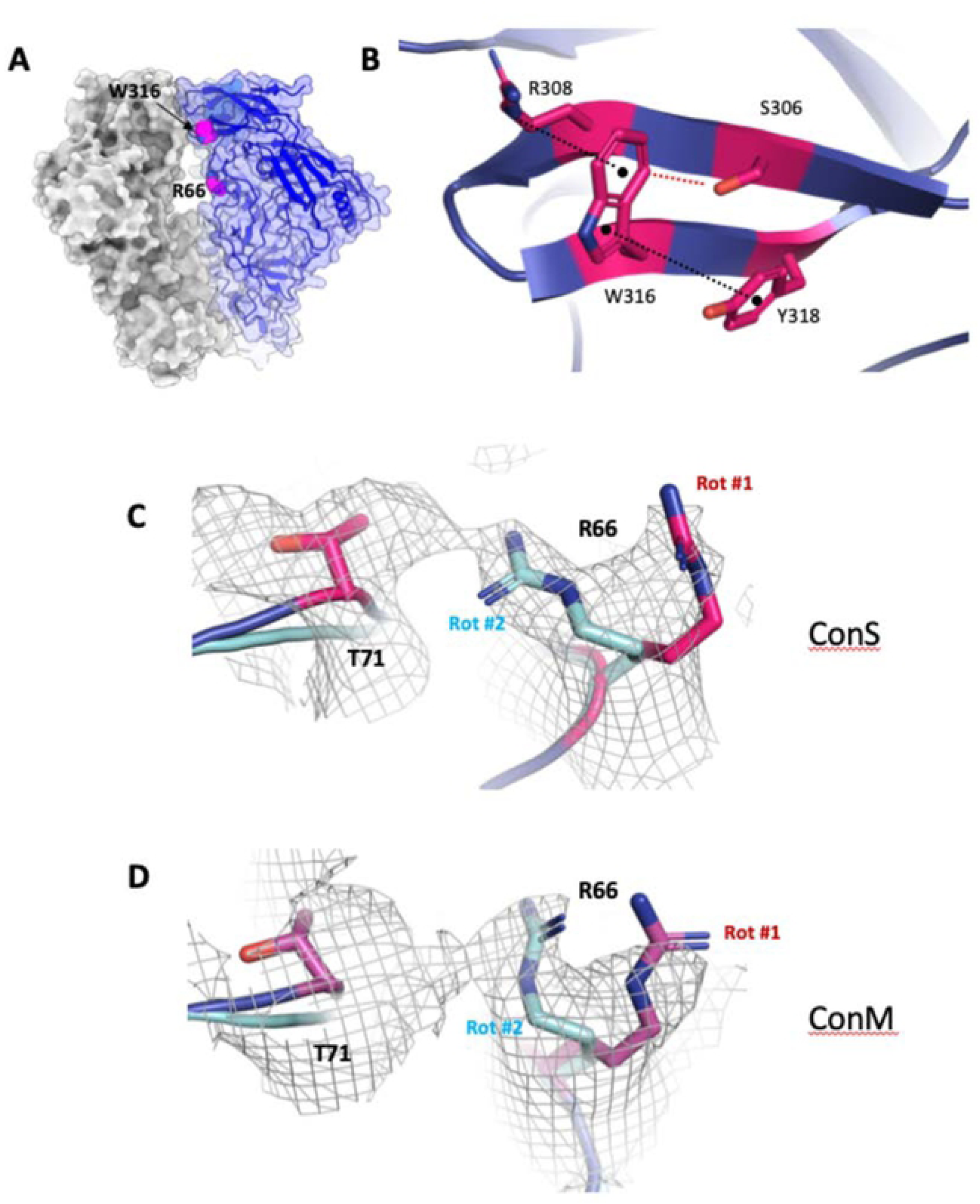
Structural details of stabilizing mutations A316W and H66R present in Cons and ConM. (**A**) Surface representation of Cons structure, with ribbon representation of one protomer shown. The W316 and R66 side chains are shown as magenta spheres. (**B**) Zoomed in view of W316 and the base of the V3 loop in the Cons structure, highlighting a hydrophobic stacking interaction with Y318, a cation-pi interaction with R308, and a possible weak H-bond with S306. (**C**) Structure and cryo-EM density of two possible rotamers for the R66 side chain in Cons. R66 is modelled as rotamer #1 in both deposited Cons and ConM cryo-EM structures, but rotamer #2 is equally likely. In this conformation, R66 would probably form stabilizing contacts with T71. (**D**) Same as in (**C**), but for ConM cryo-EM structure.

**Supplementary Figure 11.**
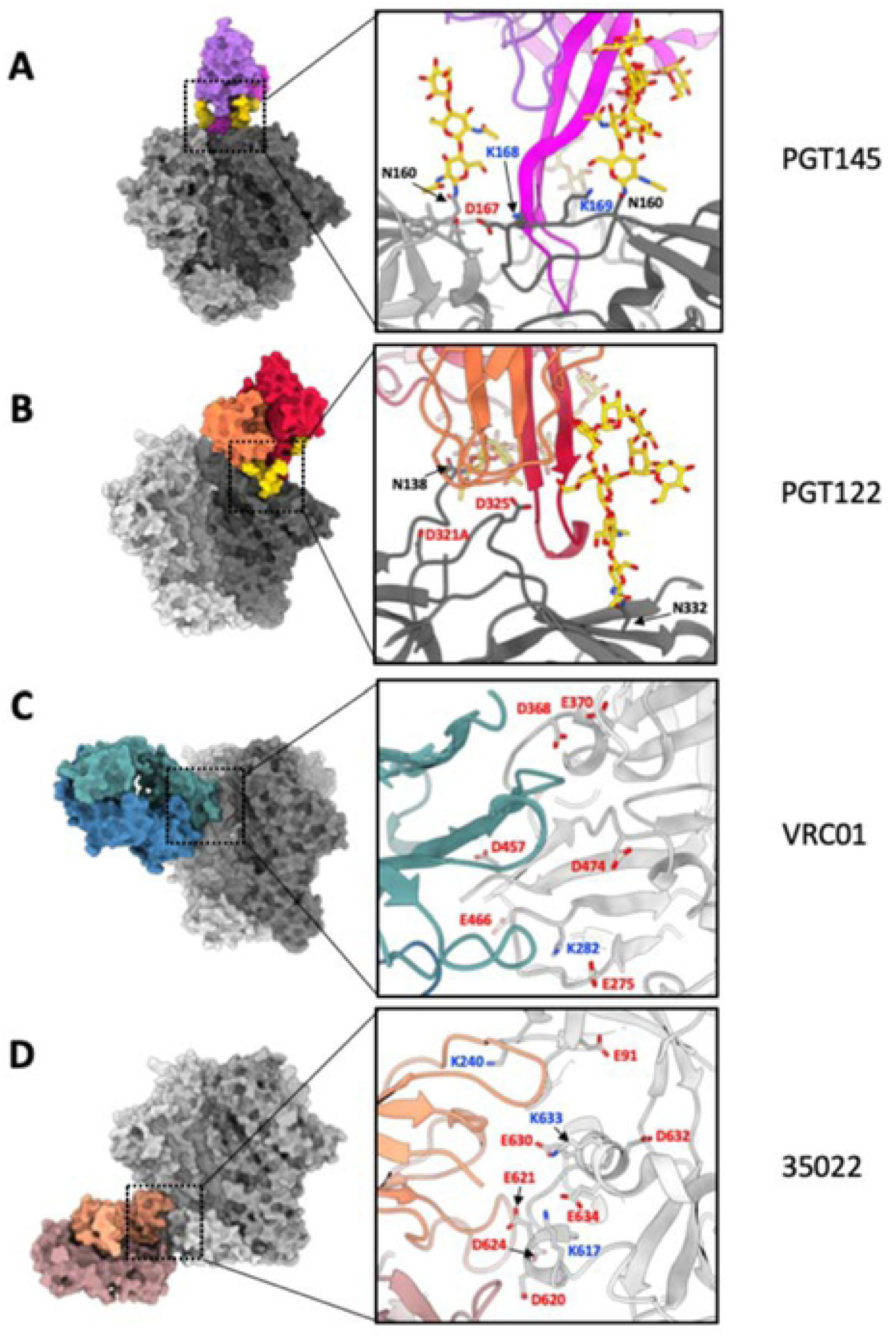
Structures of select bnAb eptiopes and exposure of glutamate, aspartate, and lysine side chains. (**A**) BG505-PGT145 cryo-EM structure (PDB 5V8l). PGT145 binding is to a large extent driven by glycan contacts. (**B**) ConS­ PGT122 cryo-EM structure (this paper), showing, as with PGT145, a glycan-dependent epitope minimally by EDC. (**C**) Cons cryo-EM structure with VRC01 docked in, based on PDB 6V8X. Though not directly observed in the cryo-EM structures, a fraction of exposed Glu and Asp side chains could have been modified by EDC crosslinking, accounting for reduced binding to EDC modified Cons and ConM. Alternatively, the limited conformational flexibility in crosslinked SOSIPs could prevent rearrangements required to mediate high affinity CD4 binding site bnAb binding. (**D**) Cons cryo-EM structure with docked 35022, based on PDB 5W6D. Although 35022 binding to EDC-crosslinked Cons and ConM is largely conserved (as with VRC01) numerous exposed Glu and Asp side chains could potentially be modified and impact bnAb binding at this site.

**Supplementary Figure 12.**
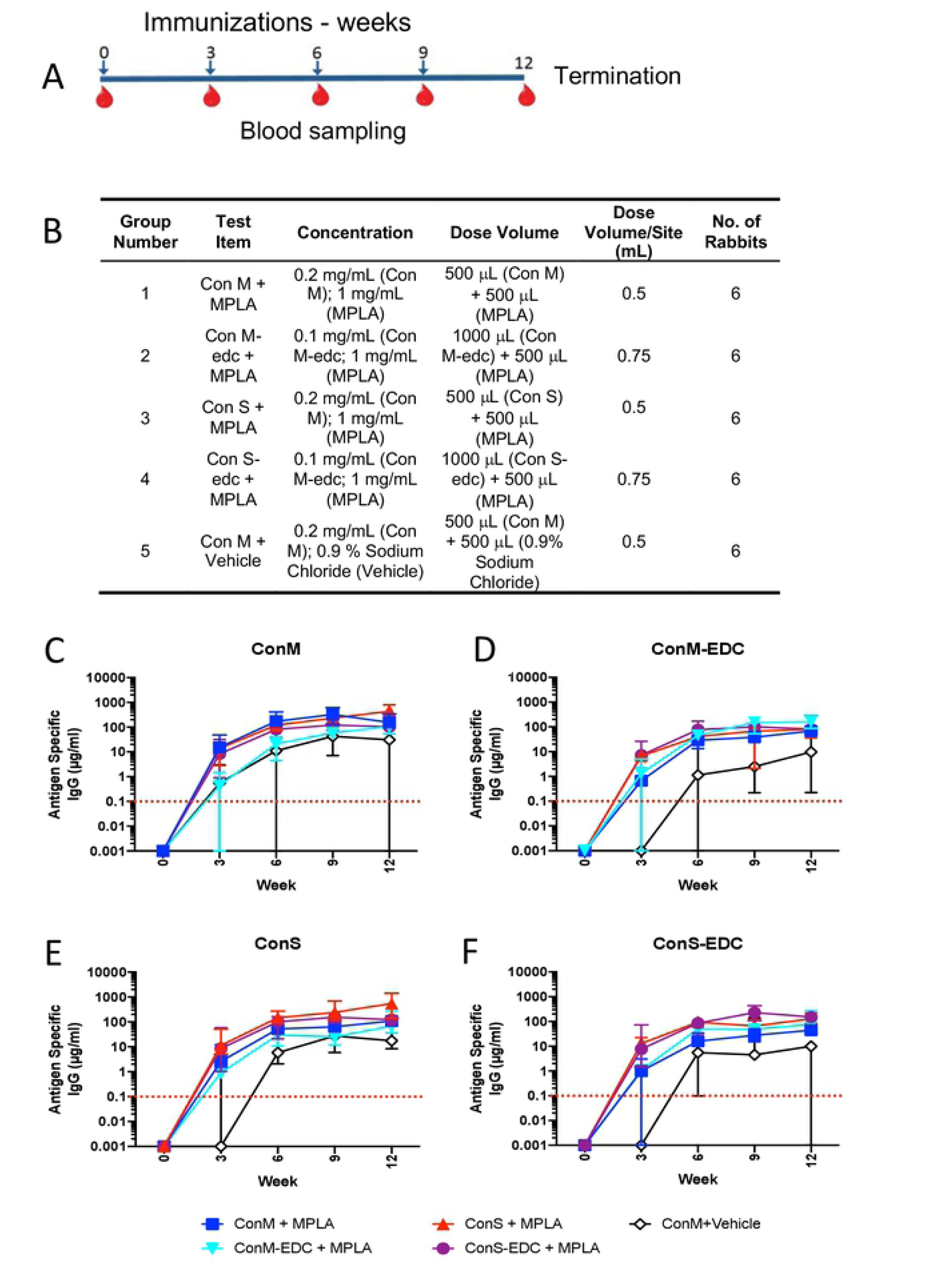
Rabbit lgG responses to antigen in MPLA immunization for toxicity study. **A**) immunization and sampling regimen where arrows represent immunizations and red drops represent bleeds. The protocol was terminated at week 12. **B**) Schedule of dosing and volumes, animals were injected at 2 sites per administration. Volumes differ between unmodified and EDC cross-linked administrations due to differing starting concentrations of GMP product. **C-F**) lgG purified from sera from rabbits immunized as in **A**) were titrated onto plates coated with: **C**) ConM, **D**) ConM-EDC, **E**) Cons, **F**) ConS-EDC,and result s reported as binding curves in µg/mL lgG.

**Supplementary table 1.**
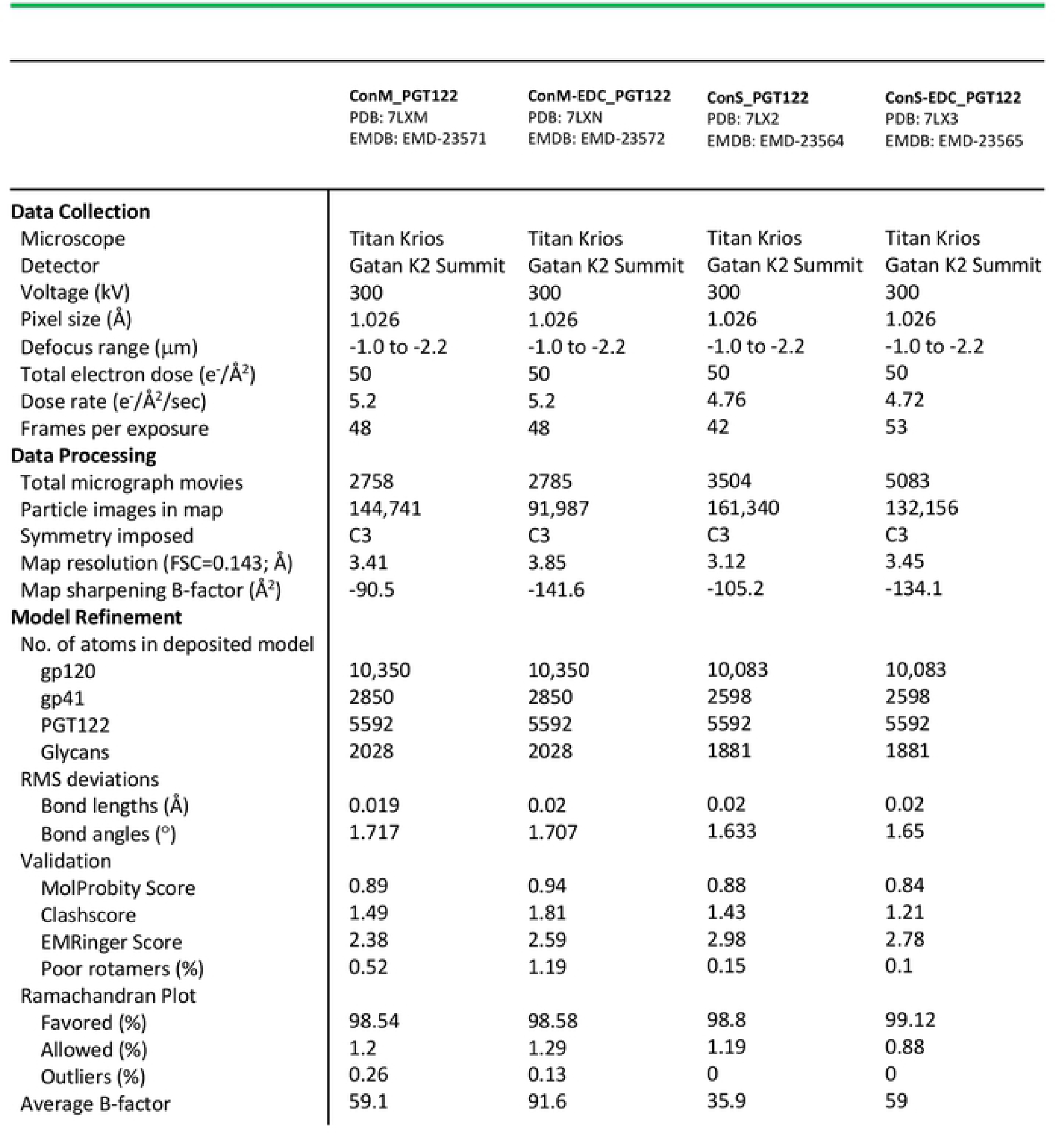
Cryo-EM data collection parameters and model-building statistics

**Supplementary Table 2.** Summary of potential and observed crosslinks in Cons and ConM. A separation distance of <6Å between Glu/Asp and Lys side chains was used to define potential crosslinks, in addition to geometric constraints as observed by the structure. See Methods for determination of observed crosslinks.

| Potential Crosslinks |  |  |  |  |  |  |  |
| --- | --- | --- | --- | --- | --- | --- | --- |
| <u>ConS</u> |  |  |  | <u>ConM</u> |  |  |  |
| <u>Intra-protomer</u> |  | <u>Inter-protomer</u> |  | <u>Intra-protomer</u> |  | <u>Inter-protomer</u> |  |
| D47 | K46 | D167.A | K169.C | D47 | K46 | D167.B | K169.E |
| D47 | K487 |  |  | D47 | K487 | E654.B | K601.D |
| D83 | K227 |  |  | E64 | K207 | E659.B. | K601.D |
| E91 | K487 |  |  | E83 | K227 |  |  |
| D107 | K574 |  |  | E91 | K487 |  |  |
| D113 | K117 |  |  | D107 | K574 |  |  |
| D167 | K168 |  |  | D113 | K117 |  |  |
| D180 | K421 |  |  | D133 | K155 |  |  |
| D230 | K232 |  |  | E153 | K178 |  |  |
| D230 | K240 |  |  | D167 | K168 |  |  |
| E267 | K231 |  |  | D180 | K421 |  |  |
| E268 | K231 |  |  | D230 | K232 |  |  |
| E268 | K232 |  |  | D230 | K240 |  |  |
| E269 | K232 |  |  | E267 | K231 |  |  |
| E269 | K348 |  |  | E268 | K231 |  |  |
| E275 | K282 |  |  | E268 | K232 |  |  |
| E275 | K97 |  |  | E269 | K232 |  |  |
| E351 | K347 |  |  | E269 | K348 |  |  |
| E351 | K348 |  |  | E275 | K282 |  |  |
| E370 | K421 |  |  | E275 | K97 |  |  |
| E466 | K356 |  |  | E290 | K340 |  |  |
| E492 | K46 |  |  | E351 | K347 |  |  |
| E492 | K490 |  |  | E351 | K348 |  |  |
| E621 | K617 |  |  | E370 | K421 |  |  |
| E630 | K633 |  |  | E466 | K356 |  |  |
| D632 | K46 |  |  | E492 | K46 |  |  |
| E634 | K617 |  |  | E492 | K490 |  |  |
|  |  |  |  | E584 | K588 |  |  |
|  |  |  |  | E621 | K617 |  |  |
|  |  |  |  | E630 | K633 |  |  |
|  |  |  |  | D632 | K46 |  |  |
|  |  |  |  | E634 | K617 |  |  |
| Observed Crosslinks (all intra-protomer) |  |  |  |  |  |  |  |
| <u>ConS</u> |  |  |  | <u>ConM</u> |  |  |  |
| E268 | K232 |  |  | E267 | K231 |  |  |
| E269 | K348 |  |  | E268 | K232 |  |  |
| E293 | K337 |  |  | E290 | K340 |  |  |
| E351 | K348 |  |  | E485 | K231 |  |  |
| E632 | K46 |  |  |  |  |  |  |
| E634 | K617 |  |  |  |  |  |  |

